# Fully Modified SpyCas9 Guide RNAs Enable Robust Genome Editing In Cells and In Vivo

**DOI:** 10.64898/2026.05.28.725424

**Authors:** Kim Anh Vu, Han Zhang, Nadia Amrani, Gitali Devi, Nicholas Gaston, Jonathan Lee, Stacy A. Maitland, Zexiang Chen, Dimas Echeverria, Pengpeng Liu, Karthikeyan Ponnienselvan, Matthew B. Hanlon, Connor Lucas, Kevin Luk, Jacquelyn Sousa, David Cooper, Alyxandr Srnka, Julia M. Rembetsy-Brown, Aditya Valji Ansodaria, Nathan Bamidele, Aamir Mir, Ken Yamada, Julia F. Alterman, Anastasia Khvorova, Scot A. Wolfe, Erik J. Sontheimer, Jonathan K. Watts

## Abstract

Precision engineering of CRISPR/Cas9 components has advanced genome editing toward therapeutic applications. Completely chemically stabilized guide RNAs (gRNAs) have the potential to improve *in vivo* editing efficacy while enabling greater flexibility in delivery strategies. However, previous generations of fully modified guides have been associated with reduced Cas9 activity. Here, we employed an iterative, structure-guided optimization strategy to systematically introduce chemical modifications at each position of SpyCas9 gRNAs. Extending beyond commonly used nucleotide modifications, we incorporated 2’-amino-RNA, 4’-thio-RNA, and extended nucleic acid (exNA) to generate gRNA designs in which 90-100% of the nucleotides are sugar- or backbone-modified. Although certain modification patterns exhibit sequence-dependent variability, we have established a growing repertoire of guides that consistently maintain or enhance editing efficacy when applied both *in vitro* and *in vivo*. Collectively, our heavily and fully modified gRNAs hold potential for applications in nuclease editing, base editing, and other genome editing tools.

**Graphical abstract:** 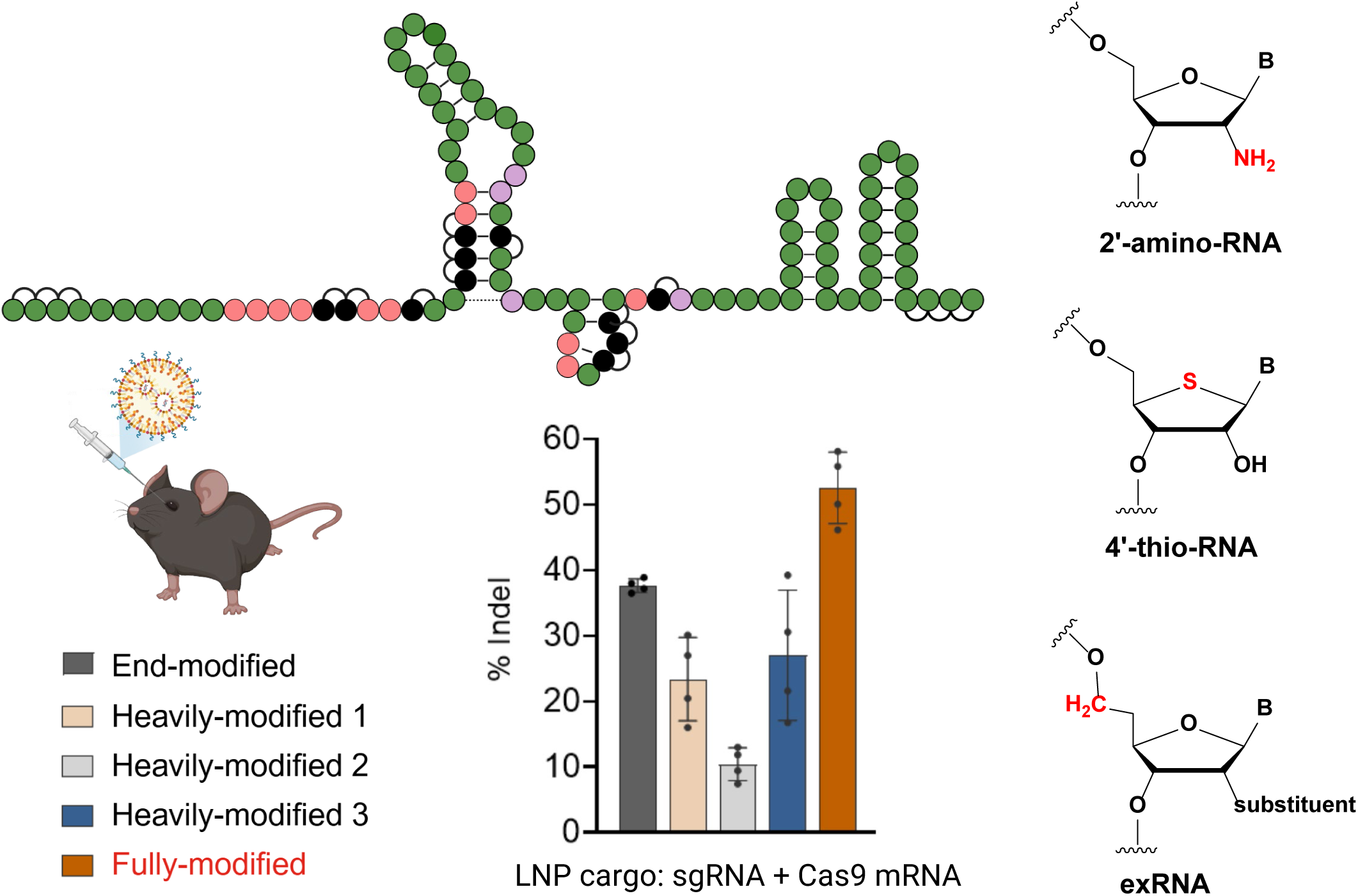

## INTRODUCTION

CRISPR/Cas9-mediated genome editing has rapidly emerged as a transformative therapeutic modality across multiple delivery routes. Rationally designed chemical modification has accelerated the clinical progress of earlier classes of RNA-based therapeutics (1–3). In CRISPR-mediated gene editing, chemically modified guide RNAs (gRNAs) enhance stability, specificity, and editing efficiency across diverse endogenous targets (4–9). However, even a single unmodified nucleotide can act as a weak link, shortening the lifetime of oligonucleotides within cells (7,10–12). Supporting this premise, a previous study showed that protecting such unmodified nucleotides by adding a modified complementary oligonucleotide to form a nuclease-resistant duplex further improves crRNA stability and potency (12). These findings suggest that the unmodified nucleotides limit gRNA durability and highlight the potential value of fully modified gRNAs to attain efficient non-viral genome editing *in vivo*.

The development of fully modified gRNAs has remained a formidable task. While our group (7) and others (4–6,8,13–15) have generated several gRNA designs that support robust Cas9-mediated editing, each design retains some unmodified nucleotides or suffers from substantially reduced activity. These hard-to-modify residues are primarily located within the repeat–anti-repeat duplex, stem-loop 1, and the linker region between stem loops 1 and 2 (see Figure 1A), where they make critical contacts with SpyCas9 through the Helical I, Bridge helix, and PAM interacting domain, respectively (16–18). Incorporating chemical modifications at these sites has been shown to reduce or even abolish editing activity (7), highlighting the structural sensitivity of these interactions and the challenge of achieving full modification without loss of function.

**Figure 1.**
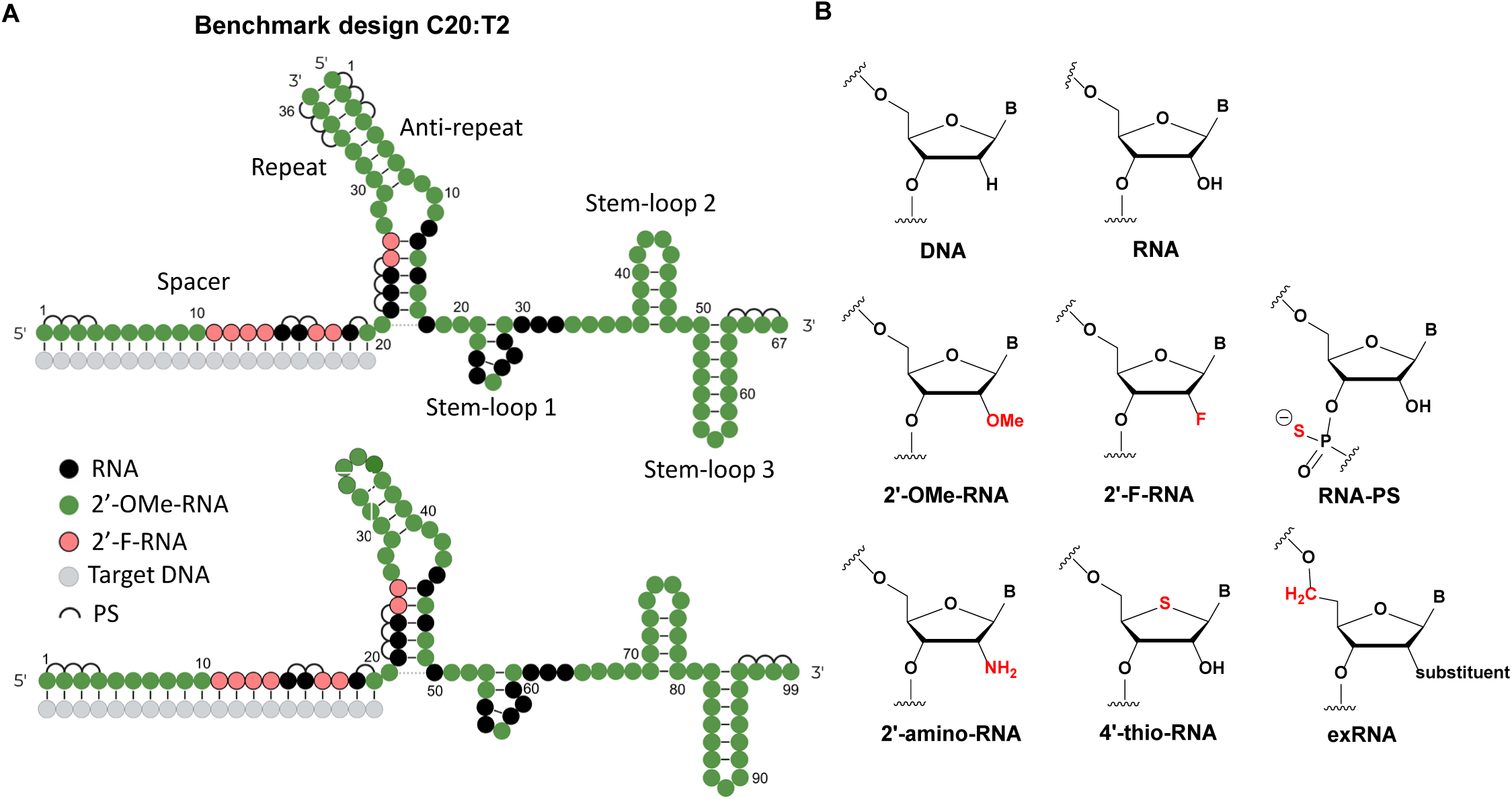
(**A**) Top: Schematic of a dual-guide RNA assembled from benchmark crRNA **C20** (C = crRNA) and tracrRNA **T2** (T = tracrRNA). Bottom: Schematic of **C20T2** single-guide RNA. RNA is shown in black; 2’-O-Me-RNA (2’-OMe) modifications in green; 2’-F-RNA (2’-F) modifications in red; and phosphorothioate (PS) linkages as half-circles. (**B**) Chemical structures of modifications used in this study. B stands for nucleobase.

This study, carried out as part of the NIH Somatic Cell Genome Editing (SCGE) Consortium (19), presents a systematic approach to develop heavily and fully modified gRNAs that maintain or enhance editing efficacy. Whereas previous approaches either focused solely on crRNAs or were largely limited to common modifications such as 2’-O-methyl-RNA (2’-OMe), 2’-fluoro-RNA (2’-F), and phosphorothioate (PS) linkages, we extended beyond these strategies by exploring additional stabilization chemistries that have not been extensively investigated in the CRISPR–Cas context. These include 2’-amino-RNA (2’-amino), which has been reported to confer moderate nuclease resistance and may partially mimic the 2’-hydroxyl group (2’-OH) through its hydrogen bonding capability (20,21). 4’-thio-RNA (4’-thio) has been shown to exhibit both endo- and exonuclease resistance, while preserving the 2’-OH substituent, potentially maintaining many of the interactions available to native RNA (22,23). Extended nucleic acids (exNA), recently developed in the context of siRNA therapeutics, have demonstrated significantly improved metabolic stability and pharmacokinetic/pharmacodynamic properties in systemic and CNS delivery (24). Accordingly, incorporation of exNA-RNA while retaining the 2′-OH might provide an effective strategy for gRNA stabilization.

Overall, through combinations of these next-generation modifications with previously developed approaches, we now present highly functional gRNAs with sugar and/or PS modifications at every position, which support efficient editing across multiple gene targets in cells and *in vivo*, in both dual-guide and single-guide RNA (sgRNA) formats, and maintain or improve activity following LNP co-delivery with mRNA relative to state-of-the-art designs.

## RESULTS

### Structure-guided refinement of crRNA

In previous work, we (7) and others (4,8,13,25) have found that 2’-OMe modifications were generally compatible across most gRNA positions, except at sites where the 2’-OH forms direct contact with SpyCas9 or where nucleotides reside in constrained binding pockets. Notably, 2’-F substitutions were tolerated at multiple sites within the critical Cas9-bound seed and repeat regions of our previously established benchmark crRNA **C20** (Figure 2A). Reasoning that 2’-F, with its small 2’-substituent, preferred C3’-endo sugar pucker, and high electronegativity, could preserve RNA-like conformation and partially mimic the electrostatic interactions that stabilize Cas9–gRNA contacts, we sought to determine whether additional 2’-F substitutions could be incorporated at the six remaining unmodified positions of **C20** (Figure 2B) without compromising guide functionality.

**Figure 2.**
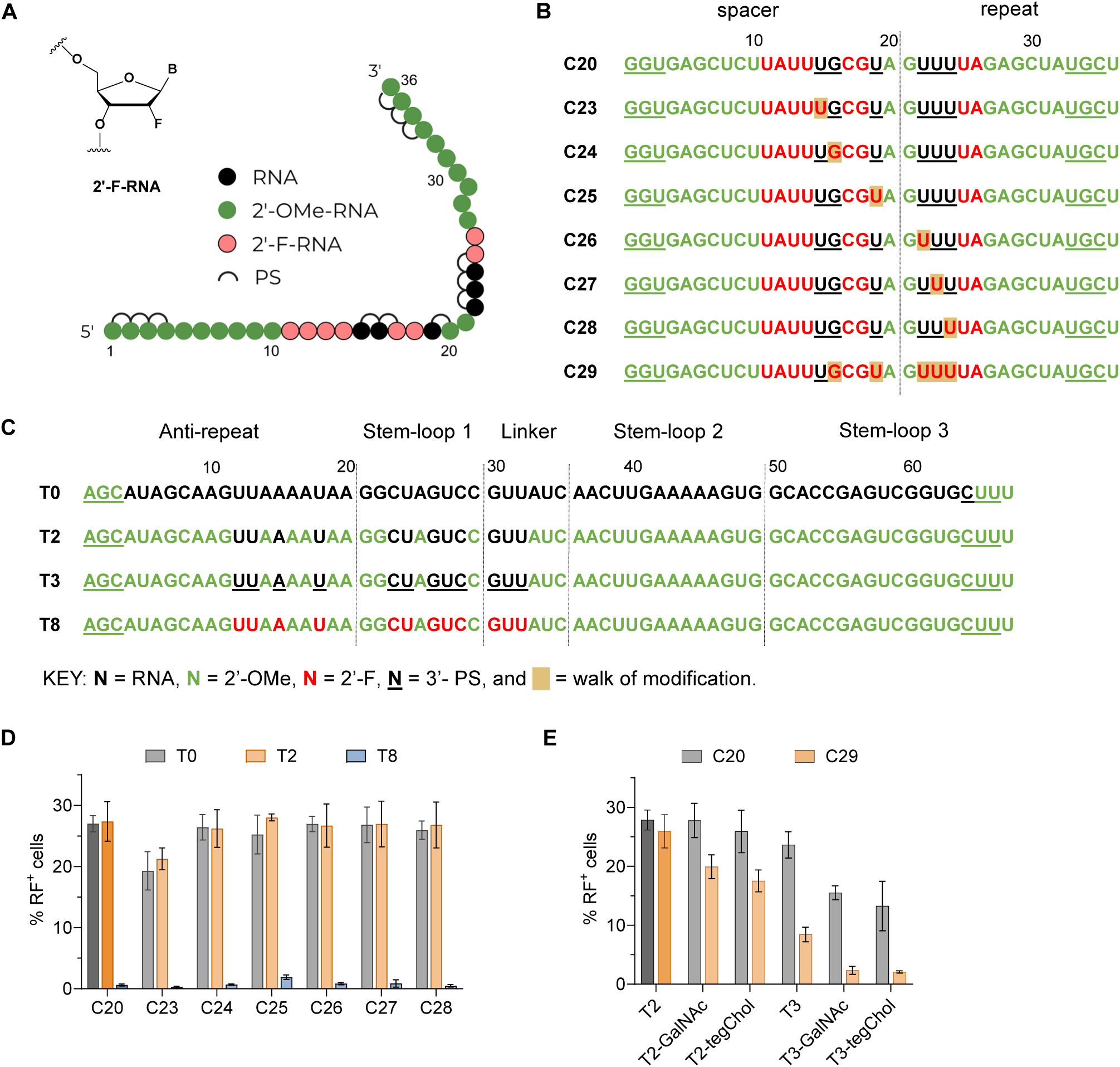
Single-position substitution of 2’-F in **C20** crRNA in HEK293T-TLR cells. (**A**) Schematic of **C20** crRNA and chemical structure of 2’-F modification. (**B**) Position-specific 2′-F substitutions introduced across individual riboses of **C20** (variants **C23**–**C29**). Substituted positions are highlighted. (**C**) Modification patterns of previously reported tracrRNAs (**T0**, **T2**, **T3**, and **T8**). (**D**) Editing efficiencies in HEK293T-TLR cells following electroporation of 20 pmol RNP assembled with crRNAs and tracrRNAs in (**B**) and (**C**). The percentage of red fluorescent-positive (RF^+^) cells quantified by flow cytometry provides an estimation of editing efficiency. Data are mean ± SD from 3 biological replicates. (**E**) Editing efficiencies of 20 pmol RNP containing **C20** or **C29** duplexed with **T2** or **T3**, with or without chemical conjugates. GalNAc, *N*-acetylgalactosamine; tegChol, tetra-ethylene-glycol-linked cholesterol. Data are mean ± SD from 3 biological replicates.

We introduced a single 2’-F substitution at each position and evaluated the resulting crRNA variants in combination with tracrRNAs **T0**, **T2**, and **T8** from our prior study (Figure 2C). We co-electroporated 25 pmol of crRNA–tracrRNA duplexes with 20 pmol of recombinant 3xNLS-SpyCas9 protein into HEK293T-TLR (Traffic Light Reporter) cells. These cells contain a GFP sequence disrupted by an insertion and followed by an out-of-frame mCherry reporter (26). In the absence of a GFP donor template, Cas9-induced double-strand breaks (DSBs) are resolved primarily through non-homologous end joining (NHEJ) to generate a spectrum of indels, some of which can restore the mCherry reading frame. Accordingly, the percentage of mCherry-positive cells reflects a subset of indel events and serves as a proxy for overall editing activity (26) (Supporting Figure S1).

Substitution with 2’-F was well tolerated at most of the remaining unmodified positions of **C20**, with only a slight reduction in activity observed for **C23** (Figure 2D). All tested crRNA variants remained compatible with **T0** and **T2**, producing editing efficiencies comparable to **C20**, whereas their activity was markedly reduced when paired with the fully modified tracrRNA **T8** (Figure 2D). This result prompted the design of **C29** (Figure 2B), in which all ribose positions in **C20** were substituted with 2’-F except for position 15. Although **C29** achieved near-saturation activity comparable to **C20** when paired with **T2**, its performance declined with other more heavily modified tracrRNAs (Figure 2E), particularly after conjugation with *N*-acetylgalactosamine (GalNAc) or tetraethylene-glycol-linked cholesterol (tegChol).

Given the variable performance of these crRNAs with fully modified tracrRNA and recognizing that Cas9–gRNA interactions depend on the context of the entire RNP complex, we next revisited other regions of our crRNA designs, seeking to improve compatibility by incorporating additional chemical variability at the previously sensitive positions within the seed and repeat regions in **C20**.

Although the SpyCas9–sgRNA–DNA crystal structure (PDB ID: 4OO8) (16) captures only a single conformational state of the complex, it provides a useful framework for predicting modifications that could negatively affect interactions between the components. Analysis of the structure revealed that spacer nucleotides at positions 4–6 form close contacts with SpyCas9 through their ribose 2’-OH and phosphate backbones within tight (<5 Å) binding pockets (Figure 3A,B). This suggests that 2’-OMe substitutions at these sites, as in **C20**, could introduce steric clashes or disrupt critical hydrogen bonds (H-bonds).

**Figure 3.**
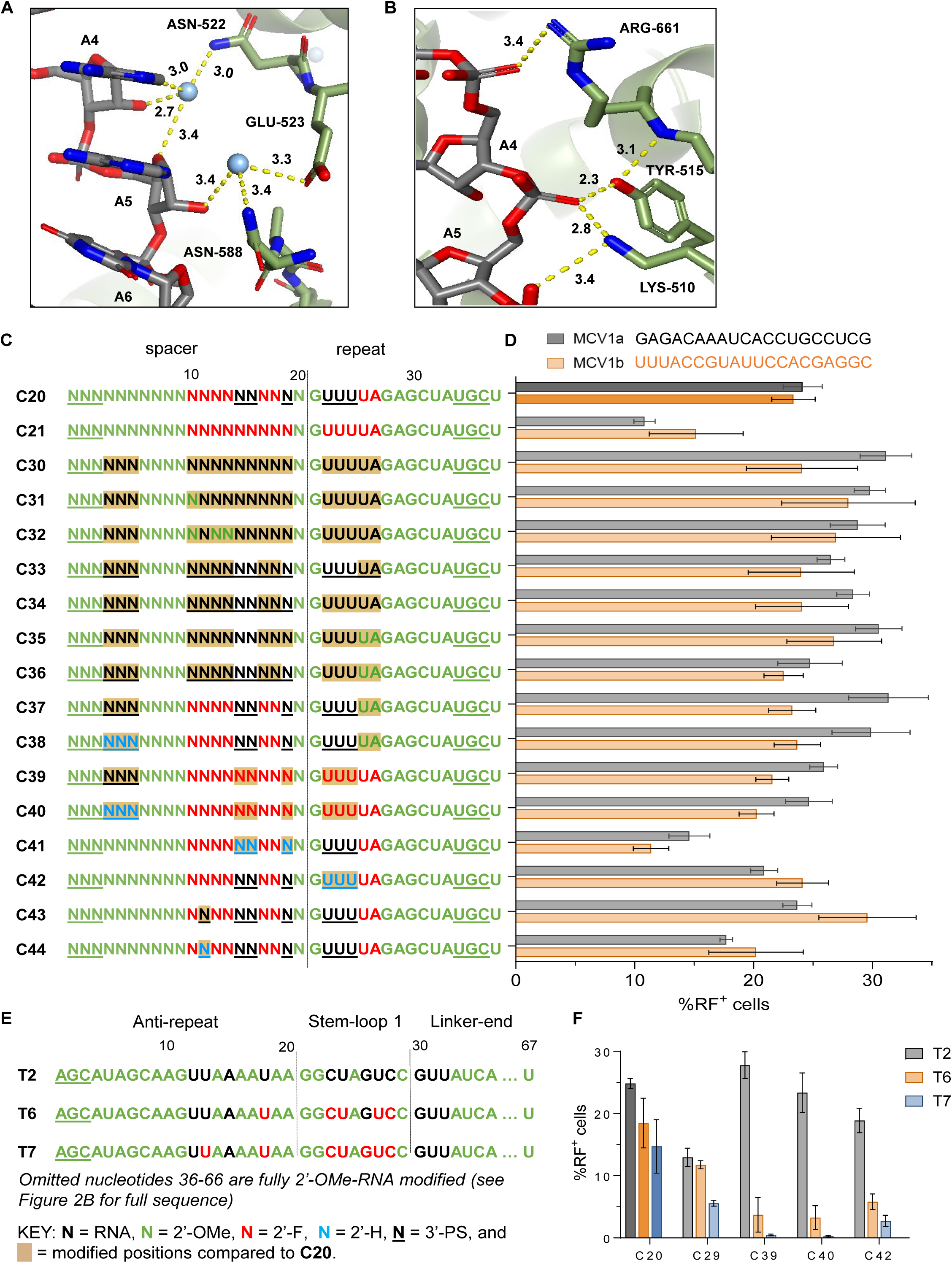
Multi-position substitution of RNA, DNA, or 2’-F with or without PS in key functional regions of C20 crRNA. (**A**) Structural model showing interactions between ribose 2’-OH groups at crRNA positions 4–5 and indicated SpyCas9 residues (PDB ID: 4OO8) (16). sgRNA, SpyCas9, and water molecules are shown in grey, green, and light blue, respectively. (**B**) Close-up of phosphate backbone contacts of crRNA positions 4–6 with indicated SpyCas9 residues. (**C**) Modification designs of previously reported (**C20** and **C21**) and newly generated crRNAs (**C30**–**C44**). (**D**) Top: Spacer sequences of 2 target loci in the HEK293T-TLR-MCV1 system. Bottom: Percentage of red fluorescent-positive (RF) cells post-electroporation of 20 pmol RNP assembled with crRNAs from (**C**) and **T2.** Data are mean ± SD from 3 biological replicates. (**E**) Schematic of previously reported tracrRNAs **T2**, **T6**, and **T7**. Omitted nucleotides 36-66 are fully 2’-OMe modified (see Figure 2C for full sequence). (**F**) Editing efficiencies in HEK293T-TLR-MCV1 cells after electroporation of 20 pmol RNP assembled using crRNAs **C20**, **C29**, **C39**, **C40**, and **C42** with **T2**, **T6**, and **T7**. Data are mean ± SD from 3 biological replicates.

To test this, we first restored native RNA at positions 4–6 and within the seed and repeat regions of **C20**, then systematically reintroduced 2’-OMe, PS, 2’-F, and DNA modifications to identify designs compatible with site-specific nuclease activity (Figure 3C). Each crRNA was paired with **T2** and tested as a 20 pmol RNP in HEK293T-TLR-MCV cells expressing the multi-Cas variant 1 Traffic Light Reporter (27,28), targeting two distinct sequence regions (MCV1a and MCV1b) within the same reporter locus.

Most crRNA designs in this panel (**C30**–**C44**) exhibited editing efficiencies comparable to **C20** for both target sites. Notably, **C39** and **C40** maintained **C20**-like activity while incorporating a higher degree of chemical modification (Figure 3C, D). Conjugation of GalNAc to either the crRNA or tracrRNA moiety of these duplexes achieved similar levels of activity to the non-conjugated guides (Supporting Figure S2). By contrast, designs that retained 2’-OMe at positions 4–6, together with minor RNA or DNA substitutions in the seed region, exhibited compromised activity (**C41**–**C44**). Although **C39** and **C40** showed improved compatibility with **T2**, **C20** remained the most consistent at generating higher editing rates with other tracrRNAs, including the heavily modified **T6** and **T7** (Figure 3E, F). Because crRNAs targeting MCV1a generally yielded higher editing activity than those targeting MCV1b, we used this target site for subsequent studies.

### Mapping functionally tolerated modifications in tracrRNA

Considering that crRNA and tracrRNA function cooperatively within the RNP complex, we next focused on optimizing the chemical modifications within the tracrRNA. Previous findings (7) identified 12 critical nucleotides within the tracrRNA, as represented in **T2**, where chemical modification significantly reduced activity; accordingly, **T2** was used as a benchmark (Figure 4A). We next generated 36 tracrRNA variants using a single-position walk strategy, introducing 2’-F or 2’-OMe sugar modifications or PS linkages at each of the 12 unmodified riboses in **T2** (Figure 4B,C). These variants were evaluated in HEK293T-TLR-MCV1 cells with the fully modified **C40** under guide-limiting conditions (*i.e.*, 2 pmol of RNP).

**Figure 4.**
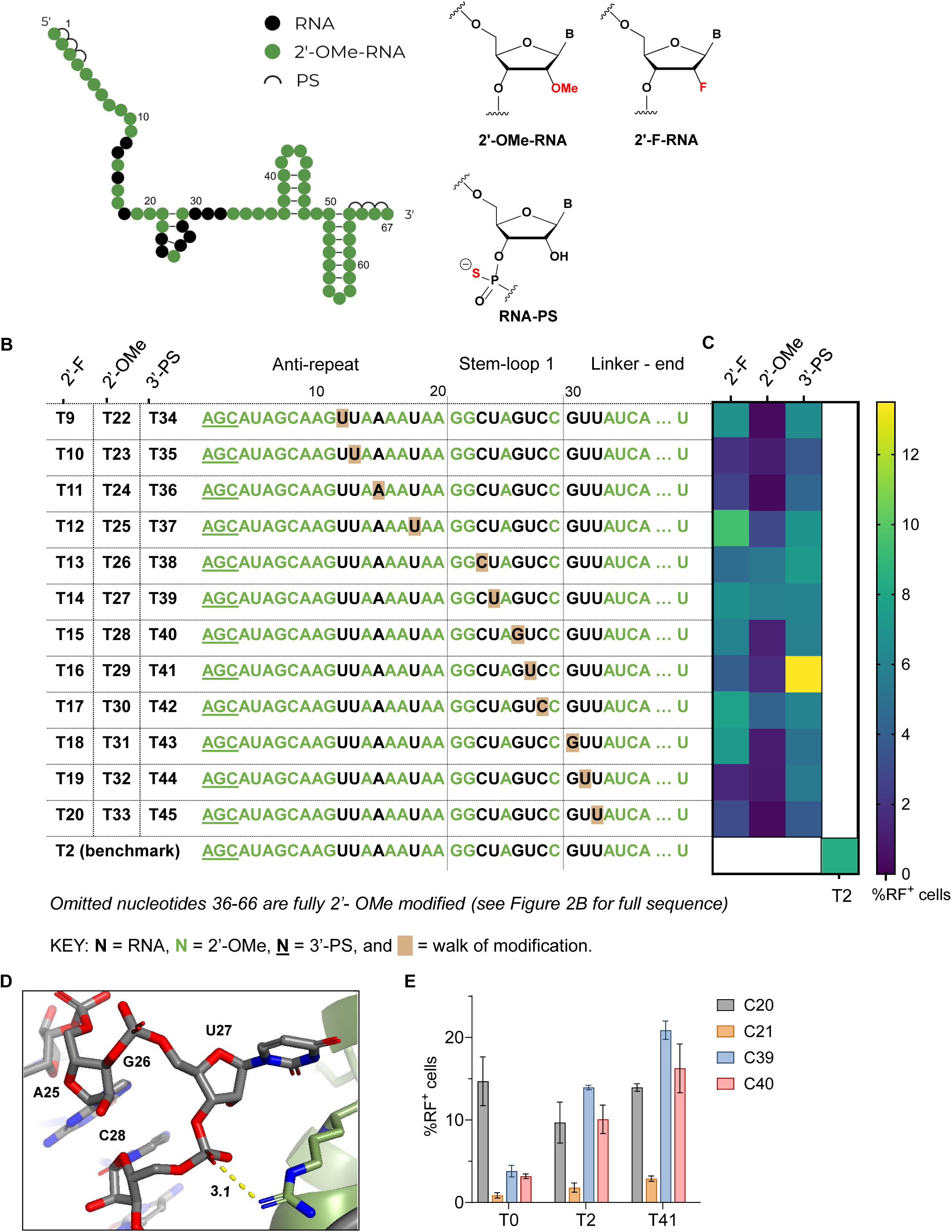
(**A**) Schematic of **T2** tracrRNA and chemical structure of 2’-F, 2’-OMe, and PS modifications. (**B**) Position-specific substitution in **T2** tracrRNA using 2’-F (**T9**-**T20**), 2’-OMe (**T22**-**T33**), or PS (**T34**-**T45**). Omitted nucleotides 36-66 are fully 2’-OMe modified (see Figure 2B for full sequence). (**C**) Heatmap representing percentage of red fluorescence-positive (RF) cells following electroporation of 2 pmol RNP assembled with **C40** crRNAs targeting MCV1a (see Figure 3B for **C40** design) and tracrRNAs from panel A. Data are mean ± SD from 3 biological replicates. (**D**) Structural context of tracrRNA U27 (corresponding to U59 in the sgRNA, PDB ID: 4OO8) (16). The sgRNA is shown in gray and SpyCas9 in green. (**E**) Percentage of RF cells after electroporation of 2 pmol RNP assembled with crRNA–tracrRNA combinations targeting MCV1a (see Figure 2B and 3C for crRNA designs). Data represent mean ± SD from 3 biological replicates.

The single 2’-F substitutions within **T2** were generally well tolerated, except at positions U13, A15, U31, and U32. Structural analysis (PDB ID: 4OO8) suggested that U13 and A15 primarily contact the recognition (REC1) domain, whereas U31 and U32 engage residues in the RuvC nuclease domain, the bridge helix, and the PAM-interacting domain, largely through phosphate backbone and 2’-OH-mediated interactions (Supporting Figure S3A). Notably, U31 adopts a C2’-endo (DNA-like) pucker, which is potentially incompatible with the strongly C3’-endo/RNA-like geometry of 2’-F modification. In contrast, single 2’-OMe substitution within **T2** almost completely abolished editing, except at positions C23, U24, and C28, where the ribose 2’-OH does not engage in H-bonds or other close interactions with SpyCas9 residues.

The single PS substitutions were more favorable than either 2’-F or 2’-OMe at nearly all ribose positions, yielding editing comparable to or higher than the benchmark **T2** (Figure 4C). Strikingly, a single PS modification at U27 (variant **T41**) nearly doubled the editing activity relative to **T2**. U27 flips out from stem-loop 2 and engages with Arg74 through electrostatic and hydrogen bonding interactions via its 3’-phosphate group (PO) (Figure 4D). The greater polarizability and charge of the PS linkage relative to PO (29,30) may strengthen this interaction with positively charged SpyCas9 residues and improve local stabilization of the RNP complex. Consistent with this, **T41** exhibited broad compatibility with most of our crRNAs (Figure 4E), enhancing editing across multiple targets, as shown below.

Having identified the effects of single-position 2’-OMe, 2’-F, and PS substitutions in T2, we next examined how these modifications performed when introduced at multiple positions simultaneously. DNA substitution was also included in this panel. We systematically introduced the same modification at two to six of the twelve unmodified nucleotides in **T2** (Figure 5A).

**Figure 5.**
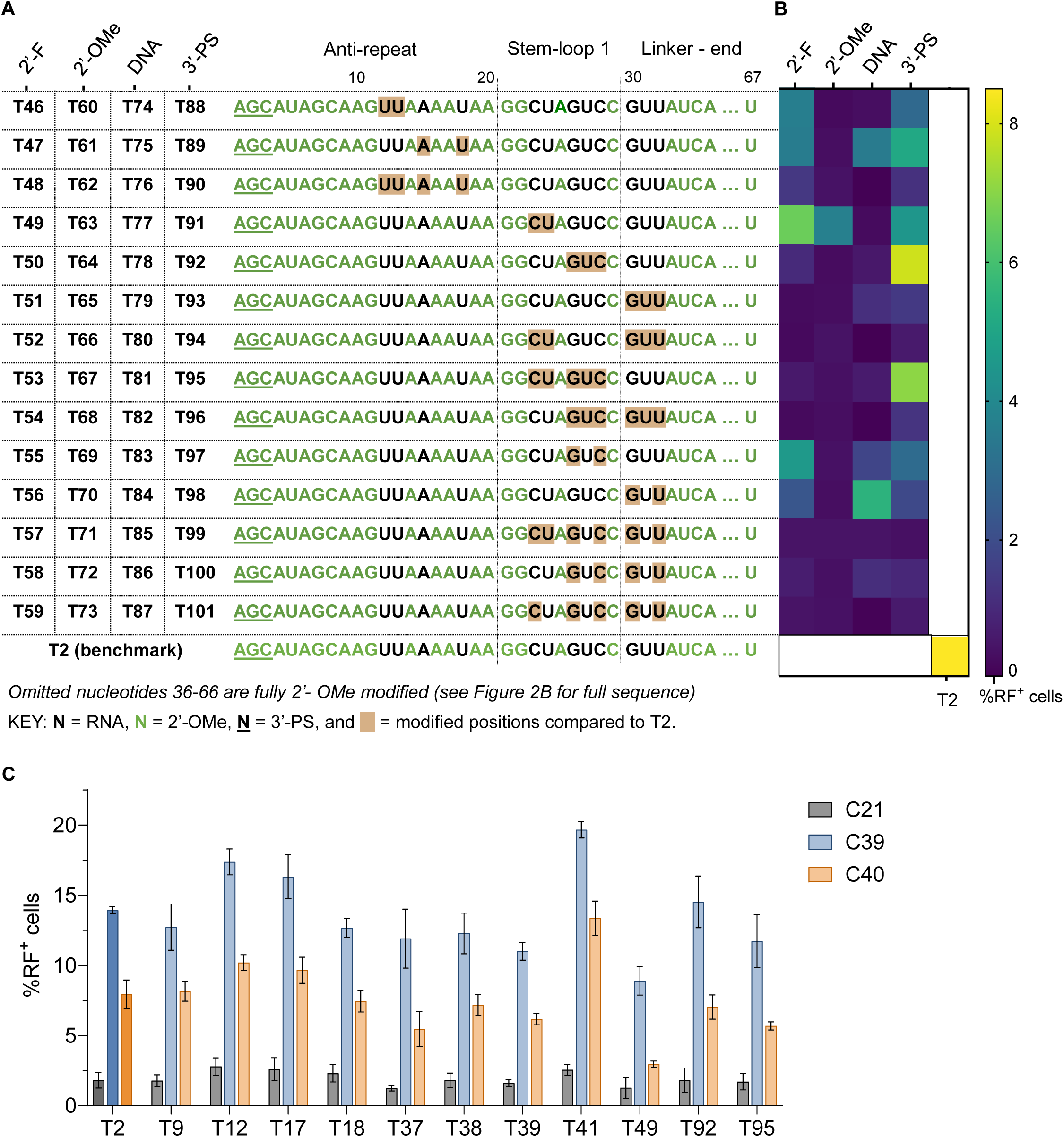
Multi-position optimization of tracrRNAs. (**A**) Multiple modifications introduced into **T2** tracrRNA using 2’-F (**T46** -**T59**), 2’-OMe (**T60** -**T73**), DNA (**T74** -**T87**), or PS (**T88** -**T101**). Omitted nucleotides 36-66 are fully 2’-OMe modified (see Figure 2B for full sequence). (**B**) Percentage of red fluorescent-positive (RF^+^) cells after electroporation of 2 pmol RNP assembled with **C40** crRNAs targeting MCV1a (see Figure 3A) and tracrRNAs from panel A. Data are mean ± SD of 3 biological replicates. (**C**) Editing efficiency of 2 pmol RNP assembled with modified crRNAs (**C21**, **C39**, and **C40**) and top-performing tracrRNAs from Figures 4 and 5. Data represent mean ± SD from 3 biological replicates.

Although single substitutions often supported robust editing, extending these modifications to a larger number of ribose positions markedly reduced activity (Figure 5B). At 2 pmol RNP, multi-substitution with either 2’-OMe or DNA abolished activity in most variants, likely owing to the combined effects of disrupted hydrogen bonds, steric hindrance, and altered local and neighboring phosphodiester backbone conformation.

Among 56 designs, only **T49**, **T92**, and **T95** retained near-**T2** editing efficiency (Figure 5B). We propose that **T49**, carrying 2’-F substitutions at C23 and U24, remained active because these positions do not form 2’-OH-mediated contacts with SpyCas9 (Supporting Figure S3B); the slight reduction in activity may reflect increased conformational constraint, as 2’-F favors a C3’-endo sugar pucker rather than the native C2’-endo at U24.

**T92** and **T95** both contain the same PS modification at U27 that enhanced activity in the context of **T41**; however, extending this modification to two or four additional neighboring linkages reduced activity. These neighboring positions are likely more sensitive to the stereochemical configuration of PS modification than U27, with only one stereoisomer (Rp or Sp) being compatible with optimal Cas9 activity, or with multiple adjacent PS linkages perturbing the local RNA conformation required for efficient Cas9 function.

To assess compatibility with the fully modified crRNAs, we tested the most active tracrRNA variants from the single- and multi-position substitution screens (Figures 4–5) with the previously reported **C21** and the optimized **C39** and **C40** from Figure 3. At 2 pmol RNP, **T12**, **T17**, **T92**, and **T95** showed activity comparable to or greater than that of the benchmark **T2. T41** remained the most active tracrRNA across all crRNA contexts, highlighting its strong activity and broad compatibility (Figure 5C).

### Single- and multi-PS substitution in crRNAs

In other classes of oligonucleotide therapeutics, the protein-binding properties of PS linkages have enabled vehicle-free delivery to multiple cell types and tissues (29,31,32). Given the increased activity observed with a single additional PS linkage in **T41**, we next investigated whether increasing PS content in the crRNA, in combination with existing 2’-OMe and/or 2’-F modifications, would also be tolerated.

Single-position PS substitutions were introduced across **C40** (Figure 6A), focusing on positions where we had limited prior experience with PS substitutions. All variants were tested in HEK293T-TLR-MCV1 cells at 2 pmol RNP. Variants **C64**-**C70** were the most active (Figure 6A, B), indicating that additional PS substitutions were best tolerated at positions near the 3’-end of the repeat region. We also evaluated a series of multi-position PS substitutions within **C40** (Figure 6C), combining well-tolerated single-position substitutions, groups that had proven to be compatible with PS in previous designs (7), and PS-modified stretches of increasing length from both ends of the crRNA. In this panel, variants **C81**-**C85** retained strong activity (Figure 6C, D), with PS substitution clustering within the repeat region. By contrast, extensive PS modification in the spacer region, as in **C79** and **C80**, was poorly tolerated.

**Figure 6.**
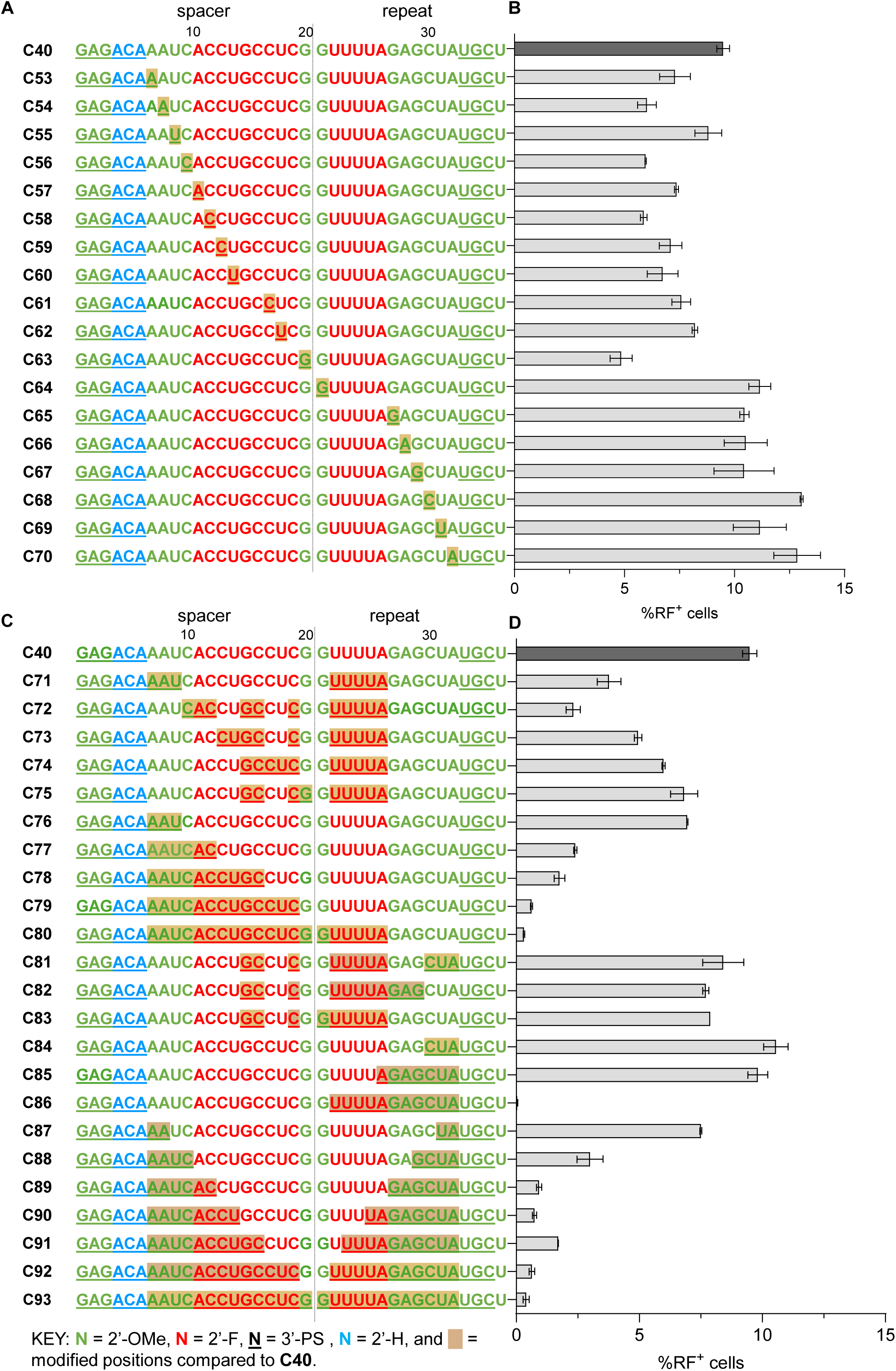
Further modification of crRNAs with PS. (**A**) Single PS substitution in crRNA **C40** targeting MCV1a reporter system. (**B**) Percentage of RF cells after electroporation of 2 pmol RNP targeting MCV1a, assembled using a modified crRNA (panel A) and **T41** tracrRNA. Data are presented as mean ± SD from 2 biological replicates. (**C**) Multi-position substitution of PS on **C40**. (**D**) Percentage of RF cells following electroporation of 2 pmol RNP targeting MCV1a, assembled using modified crRNAs (panel C) and **T41**. Data are presented as mean ± SD from 2 biological replicates.

Together, these results indicate that combinatorial substitutions of 2’-OMe, 2’-F, and PS in specific regions (**C64**-**C70** and **C81**-**C85**) can support robust editing, highlighting substantial flexibility for chemical optimization. Nonetheless, effective design requires careful consideration of both local and global structural constraints, as the impact of individual modifications remains highly context dependent.

### Optimized crRNA:tracrRNA efficiently edits endogenous targets in cells

Having identified the most active crRNA and tracrRNA variants through iterative optimization in reporter systems, we next tested their performance at endogenous genomic targets. The most active and highly modified crRNAs (**C29**, **C39**, **C40,** and **C42**) were paired with **T2** at a 5 pmol RNP dose to target the mouse *Pcsk9* (*mPcsk9*) gene in Hepa1-6 cells. The benchmark design **C20** was included in this experiment for comparison. Indel frequencies were quantified using amplicon sequencing.

Despite strong activity in the MCV1a reporter system (Figure 3B, F), **C39** and **C40** exhibited poor editing at the *mPcsk9* locus (Figure 7A). In contrast, **C29** and **C42**, which were less active in MCV1a targeting (Figure 3F), outperformed **C39** and **C40** and maintained **C20**-like activity at *mPcsk9* when delivered at 5 pmol RNP. GalNAc conjugation of either the crRNAs or **T2** did not alter activity, with conjugated guides performing comparably to their non-conjugated counterparts (Supporting Figure S4)

**Figure 7.**
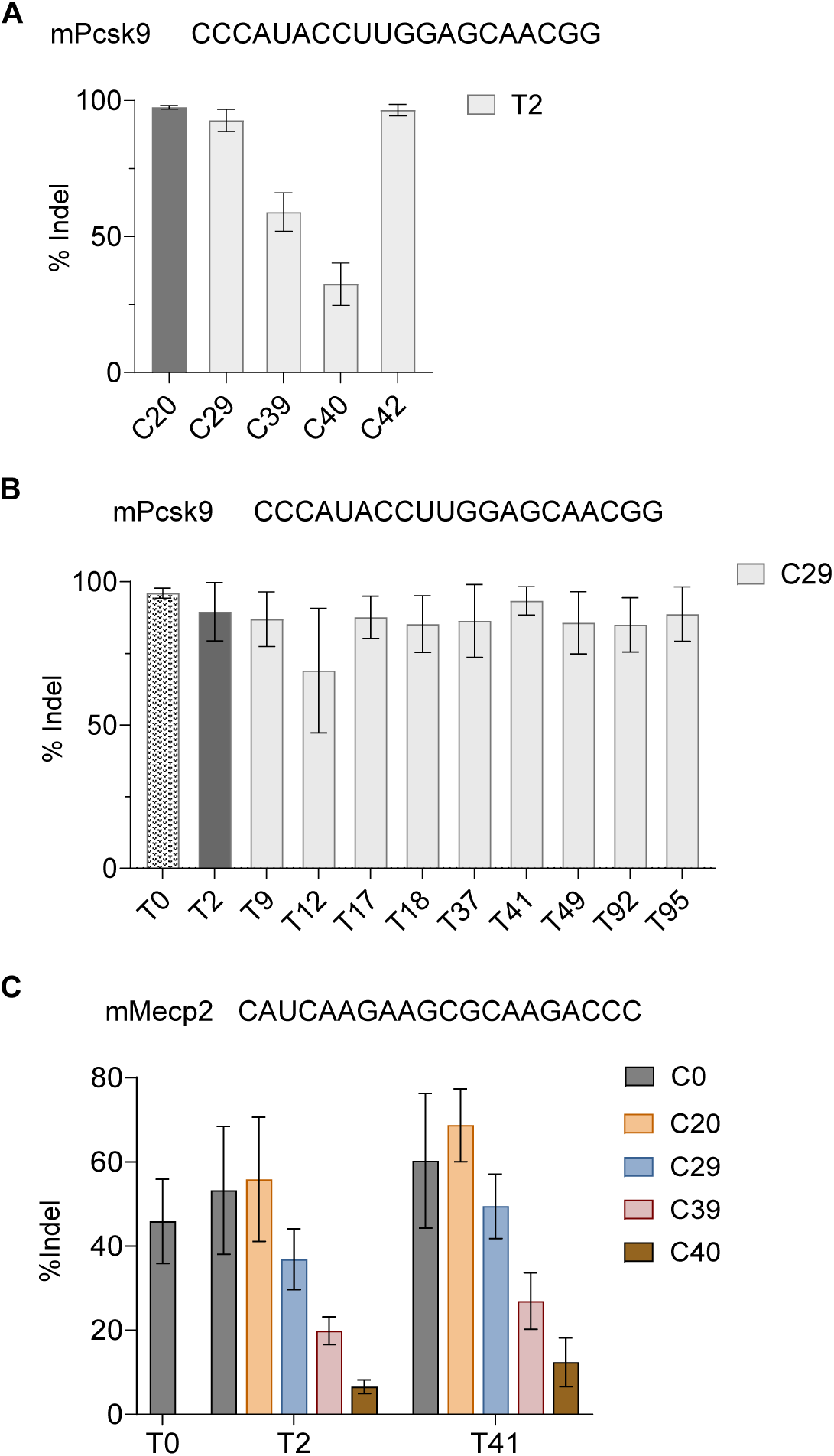
Editing of endogenous loci in cultured cells using optimized chemically modified crRNAs and tracrRNAs. (**A**) Top: Spacer sequence of *mPcsk9* targeting crRNA. Bottom: The bar graph represents the indel percentage of 5 pmol RNPs assembled using **C20**, **C29**, **C39**, **C40**, and **C42** in combination with **T2** tracrRNA in Hepa 1-6 cells. Indel percentages were quantified by amplicon sequencing. Data represent mean ± SD from 2 biological replicates. (**B**) Top: Spacer sequence of *mPcsk9* targeting crRNA. Bottom: The bar graph represents the indel percentage of 5 pmol RNPs assembled using **C29** with functional tracrRNAs **T2**, **T9**, **T12**, **T17**, **T18**, **T37**, **T41**, **T49**, **T92**, and **T95**. Data represent mean ± SD from 3 biological replicates. (**C**) Top: Spacer sequence of *mMecp2* targeting crRNA. Bottom: The bar graph represents the indel percentage of 20 pmol RNP targeting the *mMecp2* locus in MEF cells. Data are mean ± SD from 3 biological replicates.

We then paired the most active and extensively sugar-modified crRNA at this target, **C29**, with potent tracrRNA variants at the same RNP dose. Most combinations reached near-saturating activity, with only modest variability observed for **C29:T12** (Figure 7B). Collectively, these results underscore the sequence-dependent effects of modified gRNAs, particularly within the crRNA region, which contains the variable 20-base spacer sequence.

To further investigate this observation, we evaluated the same set of crRNAs with **T2** and **T41** at another endogenous locus, *mMecp2.* In primary mouse embryonic fibroblast cells, **C20** outperformed **C29**, and the **C20:T41** combination produced markedly higher editing than both the end-modified (**C0:T0**) and our benchmark (**C20:T2**) patterns. Overall, **T41** emerged as the most versatile tracrRNA, demonstrating broad compatibility with diverse crRNAs across multiple genomic targets in cells.

### Assessing the compatibility of 2’-amino, 4’-thio, and exNA modifications in crRNA and tracrRNA

After optimization with common chemical modifications, we found that although these substitutions were well tolerated at most positions in gRNAs, they could disrupt key interactions that stabilize the RNP complex, particularly H-bonds between the ribose 2’-OH group and SpyCas9 residues, resulting in reduced activity. We therefore explored additional chemistries that could enhance nuclease stability while retaining hydrogen-bonding capability, including 2’-amino and 4’-thio modifications (20–23).

Due to the complexity and cost of phosphoramidite synthesis, only 2’-amino-C, 2’-amino-U, and 4’-thio-U monomers were prepared for screening. Using the same single- and multi-position substitution strategies described above, each modification was introduced at the remaining unmodified residues in **C20** (Figure 8A). At 5 pmol RNP, most crRNA variants in this panel showed reduced activity when paired with **T2** against MCV1a, except for **C124** (4’-thio-U at U24), **C131** (2’-amino-C at C16), and **C132** (2’-amino-C at C19) (Figure 8B). In most cases, the reduced activity likely reflects conformational and/or steric effects that impair pairing either with the DNA target or with the tracrRNA anti-repeat, even when 2’-OH contacts were preserved or partially compensated.

**Figure 8.**
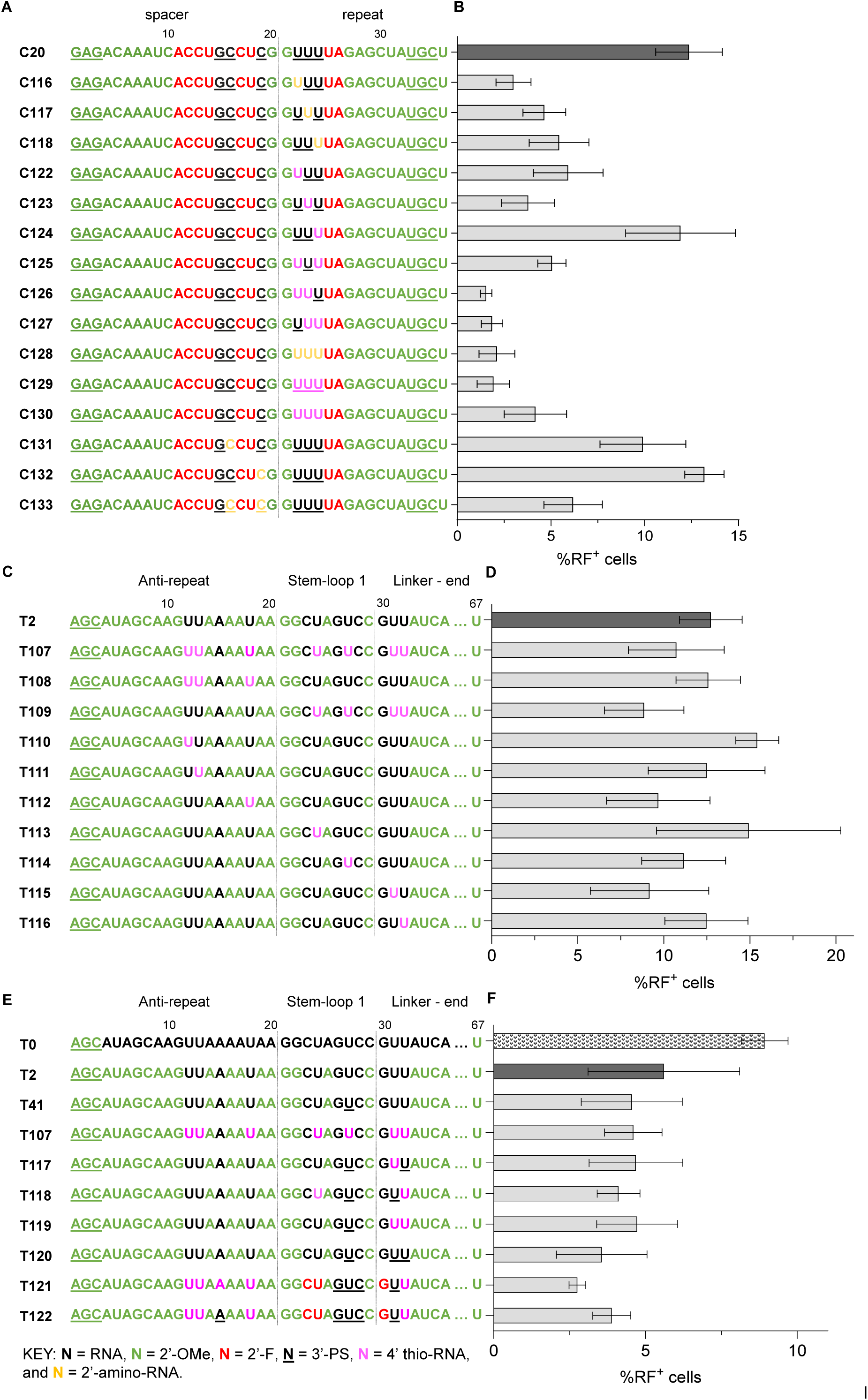
(**A**) Optimization of **C20** crRNA with modifications that maintain H-bonding substituents at the 2’-position, 2’-amino-RNA, or 4’-thio-RNA. (**B**) Percentage of RF^+^ cells upon electroporation with 5 pmol RNPs targeting MCV1a, assembled using the modified crRNAs (panel A) and **T2**. Data are presented as mean ± SD from 3 biological replicates. (**C**) Optimization of **T2** tracrRNA with 4’-thio-U. (**D**) Percentage of RF^+^ cells following electroporation of 5 pmol RNPs targeting MCV1a, assembled using the modified tracrRNAs (panel C) and **C20**. Data are presented as mean ± SD from 2 biological replicates. (**E**) Combinatorial substitutions of 2’-OMe, 2’-F, PS, and 4’-thio modifications in **T2**. (**F**) Percentage of RF^+^ cells post-electroporation from 2 pmol RNPs targeting MCV1a, assembled using the modified tracrRNAs (panel E) and **C20**. Data are presented as mean ± SD from 3 biological replicates.

Evaluation in tracrRNA was limited to 4’-thio modifications because the 2’-amino phosphoramidites showed low coupling efficiency. When paired with **C20** at 5 pmol RNP, most 4’-thio-modified tracrRNA variants (**T107**-**T116**) exhibited activity comparable to **T2** (Figure 8C, D). Notably, 4’-thio-U substitutions were generally well tolerated across all unmodified uridine positions in **T2**, as demonstrated by **T107**.

Building on these observations and our earlier screens (Figure 4-5), we designed **T117**-**T122** by combining favorable modifications at each position to generate highly modified tracrRNAs that retain activity (Figure 8E). At 2 pmol RNP, most tracrRNAs in this series showed modest reductions in activity relative to the end-modified **T0** (Figure 8F). **T121** and **T122**, each containing >92% sugar modification and PS linkages at all remaining unmodified riboses, retained ∼50% of **T0** activity at this low dose. Although reduced relative to **T0**, this level of activity is notable given the extreme extent of chemical modification and compares favorably with our previously published fully modified guides, which require a dose of 100 pmol to show meaningful activity in cells (7).

### Evaluation of extended nucleic acid modifications in crRNA and tracrRNA

We also examined extended nucleic acid (exNA), a recently developed backbone modification that enhances resistance to both endo- and exonucleases (24). The extra methylene insertion between the 5’-C and phosphate group of exNA could also increase backbone flexibility and potentially relieve conformational constraints imposed by other modifications.

Accordingly, we introduced exNA individually at each uridine position (^ex^U) in **C20** (Figure 9A) and **T2** (Figure 9C), in combination with the existing 2’OH, 2’-OMe, 2’-F, and/or PS modifications at those positions (as indicated by color in the sequences in Figure 9). The resulting crRNA and tracrRNA variants were paired with **T2** and **C20**, respectively, and tested against MCV1a at a 5 pmol RNP dose.

**Figure 9.**
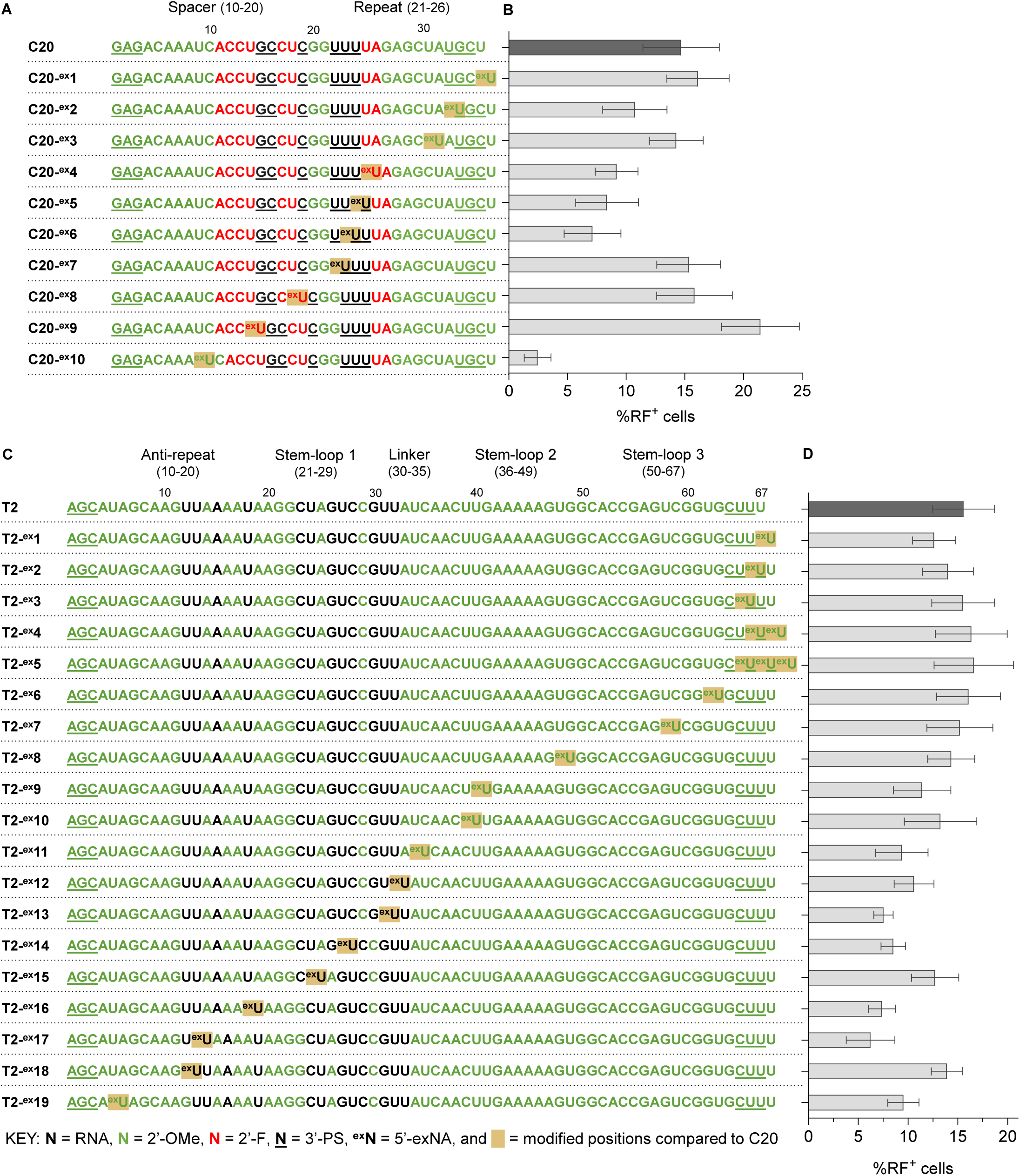
(**A**) Mapping the tolerability of exNA modification on **C20** crRNA. (**B**) Percentage of RF^+^ cells upon electroporation with 5 pmol RNPs targeting MCV1a, assembled using the modified crRNAs (panel A) and **T2** tracrRNA. Data are presented as mean ± SD from 2 biological replicates. (**C**) Optimization of **T2** tracrRNA with exNA. (**D**) Percentage of RF^+^ cells following electroporation of 5 pmol RNPs targeting MCV1a, assembled using the modified tracrRNAs (panel C) and **C20**. Data are presented as mean ± SD from 2 biological replicates. 2’-substituent is color-coded in the figure above for exNA nucleotides as well as normal nucleotides.

Although several exNA substitutions decreased activity, certain uridine positions (e.g., **C20-^ex^1**, **C20-^ex^2**, **C20-^ex^7**, **C20-^ex^8**, and **C20-^ex^9**) improved editing relative to the **C20** benchmark. Across the **T2** scaffold, exNA substitutions were generally well tolerated, with most positions maintaining activity comparable to the **T2** benchmark.

Overall, these experiments identify specific sites within both crRNA and tracrRNA that tolerate 2’-amino-RNA, 4’-thio-RNA, and exNA substitutions, expanding the repertoire of chemical modifications compatible with functional gRNAs.

### Lead crRNA and tracrRNA designs translate into highly active sgRNA in cells

Having identified lead crRNA and tracrRNA variants for MCV1a and *mPcsk9*, we next assessed their performance in sgRNA format with SpyCas9 mRNA. This step is necessary prior to *in vivo* testing via LNP delivery, because we have observed that dual guides perform poorly when co-delivered with Cas9 mRNA. Dose-response experiments were performed using 2-fold increments in sgRNA concentration. Each sgRNA targeting MCV1a or *mPcsk9* was co-electroporated with 250 ng 3xNLS-SpyCas9 mRNA. Editing efficiency increased with the sgRNA dose in a target-dependent manner, allowing a direct comparison of the lead modified designs to the end-modified **C0T0** and the published “Intellia heavily modified” design (**IHM**), which is 46% sugar-modified (8).

For MCV1a, sgRNA **C0T0** (**sgC0T0**) outperformed all of our sgRNA variants at every tested dose (Figure 10A). Although **sgC39T41** (85% sugar-modified) was the top-performing design in the dual-guide RNP format (Figure 5C), its activity as an sgRNA remained lower than **sgC0T0** and **sgIHM**. Notably, **sgC20T122**, which is 88% sugar-modified with PS substitutions at all remaining ribose positions, exhibited higher activity than C39T41 at lower doses, possibly indicating improved tolerability by Cas9. In contrast, **sgC21T41**, which is also 88% sugar-modified, was consistently less effective than the other designs.

**Figure 10.**
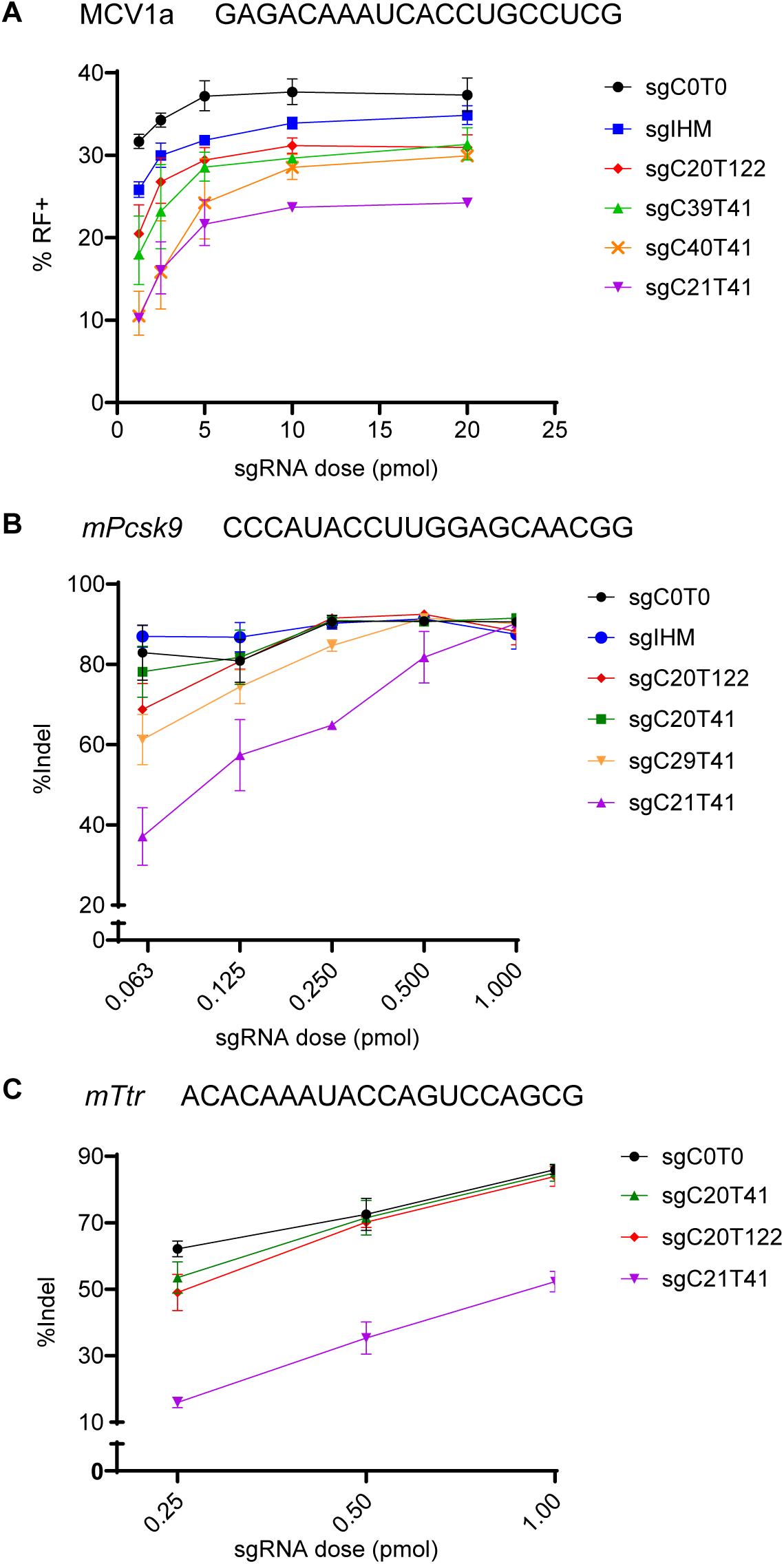
(**A–C**) Dose-response curves of lead chemically modified sgRNAs targeting (**A**) MCV1a, (**B**) *mPcsk9*, and (**C**) *mTtr* over a dose range spanning 2-fold increases in sgRNA concentration. For all panels, each sgRNA dose was co-electroporated with 250 ng of 3xNLS-SpyCas9 mRNA. Data are presented as mean ± SD from 3 biological replicates.

Across all doses, most sgRNAs supported robust editing at the *mPcsk9* locus, reaching saturation at 1 pmol sgRNA (Figure 10B). **sgC20T41** (83% sugar-modified) achieved **sgC0T0**-like activity even at the lowest dose (0.063 pmol sgRNA). Consistent with the MCV1a results, **sgC20T122** remained strongly active across the entire dose range, whereas **C21T41** exhibited a steep decline in activity with each 2-fold decrease in dose.

Excited by our results with **sgC20T122**, we wanted to validate its activity in another hepatic target, *mTtr*, to test the sequence dependence effect (Figure 10C). We included **sgC20T41** as a benchmark with consistent activity across diverse targets, **sgC20T122** as our most active fully modified sgRNA, and **sgC21T41** for comparison. Gratifyingly, both **sgC20T41** and **sgC20T122** remain highly active at the tested doses.

### *In vivo* evaluation of heavily and fully modified sgRNAs with Cas9 mRNA delivered via LNPs

To assess the functional performance of our chemically modified sgRNAs in a physiologically relevant setting, we formulated our lead sgRNA designs with 3xNLS-SpyCas9 mRNA at a 1:6 ratio in LNPs and delivered them to C57BL/6 mice via retro-orbital injections. Editing efficiency was evaluated at 2 LNP doses for each hepatic target, *mPcsk9* (0.2 and 0.8 mg/kg) and *mTtr* (0.3 and 1 mg/kg), using amplicon sequencing.

For *mPcsk9*, the fully modified **sgC20T122** outperformed the end-modified **sgC0T0** at 0.8 mg/kg and maintained comparable editing efficiency at a 4-fold lower dose, whereas all other modified sgRNA designs showed consistently reduced activity (Figure 11A). In contrast, for *mTtr*, **sgC20T41** and **sgC20T122** were slightly less active than **sgC0T0** at 1 mg/kg and exhibited a greater reduction in activity at 0.3 mg/kg (Figure 11B).

**Figure 11.**
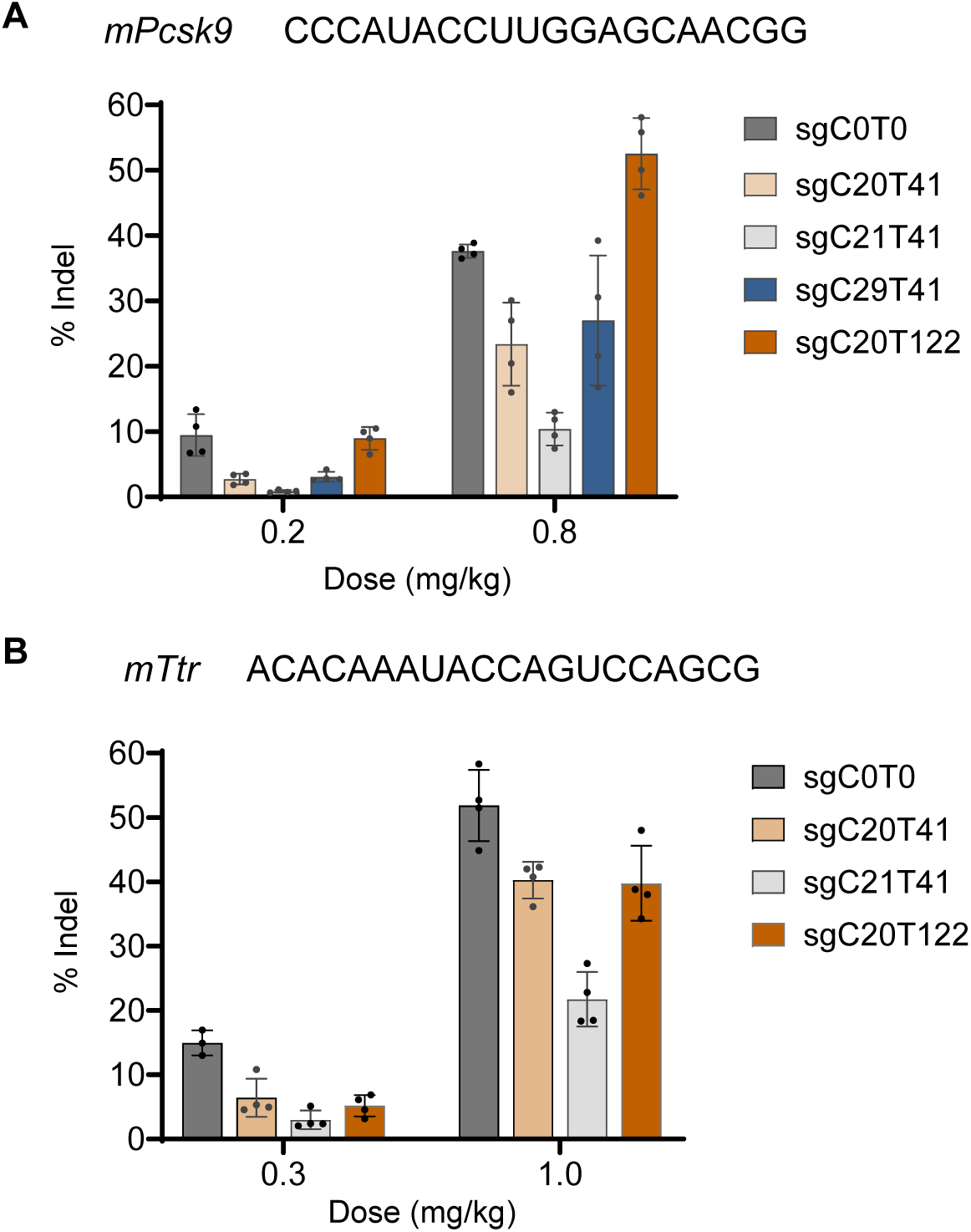
*LNP*-mediated *in vivo* genome targeting with modified sgRNAs and SpyCas9 mRNA. (**A**) *mPcsk9*. (**B**) *mTtr*. For all panels, each sgRNA was formulated with 3xNLS-SpyCas9 mRNA at a 1:6 ratio in LNPs. Data are presented as mean ± SD, n = 4.

We identified a fully modified design that supported efficient editing at the *mPcsk9* locus; however, its performance was sequence-dependent, likely due to seed-region modifications constraining guide-target base pairing and increasing sensitivity to sequence changes.

The consistent activity of **sgC0T0** across both targets suggests that guide stability is no longer the dominant limiting factor once a threshold level of chemical protection is reached. Under conditions of efficient hepatic LNP delivery, additional stabilization provides minimal benefit.

## DISCUSSION

Chemical modification of CRISPR guide RNA is a powerful strategy to improve stability, pharmacokinetics, and therapeutic potential (4–9,12–15,33). In this study, we performed a systematic, position-resolved evaluation of commonly used modifications, such as 2’-OMe, 2’-F, and PS, as well as 2’-amino, 4’-thio, and exNA, which have not been previously explored in the context of CRISPR gRNA. By combining diverse sugar and backbone chemistries, we enabled modification of regions that had previously been hard to modify without significant loss of activity. In this way, we generated highly modified (90-100%) guide designs that retain robust activity.

Our cellular data demonstrates improved editing efficiencies and increased levels of chemical modifications in this new generation of guides compared to the previously reported designs (7). The effects of individual modifications were highly context-dependent, with activity strongly influenced by local nucleotide environments and positional constraints. As such, we observed sequence dependence in the modification patterns tolerated within cRNAs. We nevertheless observed that the most active modified tracrRNAs were consistently compatible even when paired with different crRNA sequences.

As noted above, our structure-based modified gRNAs relied heavily on a single static structure model (16), which may limit the broader applicability of these modified guides given the highly dynamic nature of Cas9. A recent study has combined cryo-EM analyses and molecular dynamics simulations to investigate the effects of specific crRNA modifications across multiple conformational states (25). Extending similar approaches to study the tracrRNA or sgRNA contexts may be necessary to develop chemically modified guides with maximal compatibility with Cas9 and improved potency.

Given that our modified sgRNAs achieved activity comparable to the end-modified **sgC0T0** in cellular assays, we anticipated that they might further enhance editing *in vivo*. Following LNP-mediated delivery to the liver, the fully modified **sgC20T122** showed higher editing efficiency in targeting *mPcsk9* than **sgC0T0**, demonstrating that extensive chemical modification can improve *in vivo* performance. In contrast, other designs exhibited lower activity relative to **sgC0T0** across two hepatic targets, mirroring trends observed in cellular dose-response experiments. Moreover, for a second target (*mTtr*), while the fully modified sgRNA maintained robust editing activity, it did not surpass the end-modified sgRNA in activity. These findings are consistent with prior reports showing variable effects of extensive chemical modification on liver editing efficacy (5,8,34). The liver is highly permissive of LNP uptake following systemic administration, and the rapid endosomal escape of gRNA and mRNA enabled by advanced LNP technology may minimize exposure to endosomal and intracellular nucleases. In this physiological and delivery context, guide stability may no longer be the primary limiting factor; instead, factors such as mRNA lifetime and translation rate, and guide-Cas9 and guide-target binding kinetics, may limit the overall editing efficiency. The impact of guide modification on some of these steps may be sequence-dependent.

These caveats do not diminish the potential value of fully modified gRNAs. Rather, fully and highly modified guides may offer greater advantages in settings where delivery is less efficient, nuclease exposure is prolonged (for instance, in hard-to-deliver tissues), delivery is carried out by free uptake or separate administration of guides, or multiple doses are given. Furthermore, the modification designs described here are broadly applicable beyond nuclease-mediated genome editing and may inform guide designs for base editing, prime editing, and other SpyCas9-based modalities that rely on prolonged guide stability.

In summary, this work provides a comprehensive framework for gRNA chemical engineering by defining the tolerance and functional consequences of extensive chemical modifications across both crRNA and tracrRNA. Our study expands the toolkit for rational gRNA design and informs future efforts to balance stability, activity, sequence-dependent effects, and delivery considerations in therapeutic genome editing.

## MATERIALS AND METHODS

### Oligonucleotide synthesis and deprotection

Oligonucleotides were synthesized through solid-phase synthesis using Dr. Oligo 48 (Biolytic), K&A H-8 Synthesizer (Sierra BioSystems), and an ÄKTA Oligopilot Plus 10 (Cytiva). Phosphoramidites were obtained from ChemGenes, except for the 4’-thio phosphoramidites, which were either synthesized in-house or sourced from Sapala Organics. 2′-OMe, 2′-F, and 2′-*O*-TBDMS-protected exNA-uridine phosphoramidites were synthesized in-house according to the method previously reported (23). Phosphoramidites were prepared at 0.1 M in anhydrous acetonitrile (ACN). 2’-*O*-methyl-uridine phosphoramidites were diluted in 15% dimethylformamide in anhydrous ACN to minimize crystallization on the synthesizer. 5’-(Benzylthio)-1H-tetrazole (BTT) (ChemGenes) at a concentration of 0.25M in ACN was used as an activator. 0.05 M iodine in pyridine:water (9:1) (Tedia) was used as an oxidizer, and 0.1 M DDTT (ChemGenes) was used as a sulfurizing agent. Detritylations were performed using 3% trichloroacetic acid in dichloromethane for Dr. Oligo and K&A H-8 and 3% dichloroacetic acid in toluene (ChemGenes) for ÄKTA Oligopilot Plus 10. A mixture of 50:50 of CAP A (ChemGenes, 20% n-methylimidazole in ACN) and CAP B (ChemGenes, 20% acetic anhydride and 30% 2,6-lutidine in ACN) was used as a capping reagent. Deprotection of the nucleobase-protecting group was achieved with a (1:1) cocktail of 30% aq. Ammonium hydroxide (NH_4_OH) and 40% aq. Methylamine solution for 10 min at 65°C or with NH_4_OH at room temperature for 48 h. The oligos were flash-frozen in liquid nitrogen and dried to completion under a Speedvac concentrator. Oligonucleotides containing TBDMS protecting groups were dissolved in 115 μL DMSO (Sigma-Aldrich) at 65°C. Triethylamine (60 μL) was added, followed by 75 μL of triethylamine-trihydrofluoride. The reaction mixture was then incubated at 65°C for 2.5 hours. Following desilylation, the oligonucleotides were precipitated by adding 70 μL of 3 M NaOAc and 400 μL of isopropanol, followed by incubation at -20 °C for 30 min. The samples were then centrifuged at 14000 x g for 15 min, and the resulting pellet was washed twice with 1mL of cold 70% ethanol before being resuspended in 400 μL of RNase-free water. Desalting was performed using Amicon Ultra 0.5 mL 3 kDa MWCO centrifugal filter units (Millipore, UFC5003), followed by two ethanol washes, then three washes with RNase-free water. Samples were centrifuged at 14000 x g for 15 min, subsequently recovered by elution in RNase-free water.

### Oligonucleotide purification and desalting

Purification of oligonucleotides was performed using an Agilent 1260 Infinity HPLC system with an Agilent PL-SAX 1000 Å column (150 x 7.5 mm, 8 μm). Buffer A consisted of 20mM sodium acetate with 10% ACN, and buffer B consisted of 20 mM sodium acetate, 1M sodium bromide, and 10% ACN in water.

Fractions collected from the HPLC runs were analyzed by LC-MS using an Agilent 6530 Q-TOF mass spectrometer. Mass spectrometer was performed using electrospray ionization in negative ion mode over an m/z range of 100-32000 at a scan rate of 2 spectra/s with capillary voltage of 4000 V, fragmentor voltage of 200 V, and gas temperature of 325°C.

Liquid chromatography was carried out on an AdvancedBio oligonucleotide C18 column (2.1 x 50 mm). Buffer A consisted of 100 mM 1,1,1,3,3,3-hexafluoroisopropanol (HFIP) with 9mM triethylamine (TEA) in LC-MS-grade water, and buffer B consisted of 100 mM HFIP with 9mM TEA in LC-MS-grade methanol. A linear gradient of 5-100% buffer B over 7 min was used for all oligonucleotides at 60°C, with a flow rate of 0.85 mL/min. LC peaks were monitored at 260 nm, and data were analyzed using Agilent MassHunter software.

Once purity was confirmed, fractions were combined, desalted using Amicon Ultra 0.5 mL 3 kDa MWCO centrifugal filter units, and eluted in PBS, pH 7.4. Oligonucleotide purity was re-verified by LC-MS and fragment bioanalyzer using the DNF-470-33 Small RNA assay prior to *in vivo* administration (Supporting Figure S5). Oligonucleotide concentrations were determined by measuring absorbance at 260 nm on a Nanodrop 1000. Sequences, chemical modification patterns, and molecular weights are provided in the Supporting Table 1-3 (Excel file).

### 3xNLS-SpyCas9 purification

Protein purification for 3×NLS-SpyCas9 was performed similarly to our previous approach (35). Briefly, pET21a plasmid backbone for each Cas9 protein was transformed into *E. coli* Rosetta(DE3)pLysS cells (EMD Millipore) for protein production. Following induction, with 1M IPTG cells were pelleted by centrifugation and then resuspended with Nickel-NTA buffer (100 mM TRIS + 1 M NaCl + 20 mM imidazole + 1 mM tris(2-carboxyethyl)phosphine (TCEP), pH 7.5) supplemented with HALT Protease Inhibitor Cocktail, EDTA-Free (100×) (ThermoFisher) and lysed with a M-110s Microfluidizer (Microfluidics) following the manufacturer’s instructions. Following purification on Ni-NTA column, the protein was eluted using elution buffer (50 mM TRIS, 500 mM NaCl, 500 mM imidazole, 10% glycerol, pH 7.5). The Cas9 protein was dialyzed overnight at 4°C in 50 mM HEPES, 1 M NaCl, 1 mM EDTA, 10% (w/v) glycerol, pH 7.5. Then, the protein was batch bound to loose Capto Q resin (Cytiva) and diluted with 50 mM HEPES, 10% (w/v) glycerol, pH 7.5, until conductivity reached 25 mS/cm. Next, the protein was purified by stacked ion exchange chromatography columns (Columns = a 20ml HiPrep-Q on top of a 5 ml HiTrap-SP, Buffer A = 20 mM HEPES pH 7.5 + 1 mM TCEP, Buffer B = 20 mM HEPES pH 7.5 + 1 M NaCl + 1 mM TCEP, Flow rate = 5 ml/min, CV = column volume = 5 ml). The primary protein peak from the CEC was concentrated and buffer-exchanged in an Ultra-15 Centrifugal Filter Ultracel-30K (Amicon) to a concentration around 100 mM based on absorbance at 280 nm in 20 mM HEPES, pH 7.5, 150 mM NaCl. The purified protein quality was assessed by SDS-PAGE/Coomassie staining to be >95% pure, and protein concentration was quantified with Pierce^TM^ BCA Protein Assay Kit (Thermo Fisher Scientific). Protein was stored in –80°C until further use.

### Cell culture and electroporation

The HEK293T reporter cell lines and Hepa1-6 stable cell lines were cultured in Dulbecco’s modified Eagle’s media (DMEM, Genesee Scientific 25-500) supplemented with 10% fetal bovine serum (FBS, Gibco 26140079). All cells were incubated in a 37°C incubator with 5% CO_2_.

Electroporation of ribonucleoprotein (RNP) complexes or sgRNA/Cas9 mRNA was performed using the Neon Transfection System 10 μL Kit (ThermoFisher, MPK 1096) or the Neon NxT Electroporation System 10-μL Kit (ThermoFisher, N1096). Both systems were operated using identical parameters and yielded comparable editing efficiencies.

RNP complexes were formed by mixing crRNA, tracrRNA, and 3xNLS-SpyCas9 at a 2:2:1 molar ratio and incubating in Buffer R at room temperature for 20 min. RNP complexes or sgRNA/Cas9 mRNA mixtures in Buffer R were then added at a 1:1 ratio to 100,000 cells per sample suspended in Buffer R.

Electroporation conditions were as follows: for HEK293T cells, 1150 V pulse voltage, 20 ms pulse width, 2 pulses; for Hepa1-6 cells, 1230 V pulse voltage, 20 ms pulse width, 3 pulses. Cells were then immediately transferred to 500 μL of pre-warmed culture medium in 24-well plates and cultured for 48-72 h before analysis by flow cytometry or deep sequencing.

### Flow cytometry

Cells were trypsinized, transferred into microcentrifuge tubes, and centrifuged at 300 x g for 3 min. The supernatant was aspirated, and cell pellets were resuspended in PBS with 5% FBS before being transferred into 5 mL Polystyrene round-bottom tubes (Falcon, 352235). The percentage of fluorescent-positive cells was quantified using MACSQuant VYB flow cytometer, with 20000 events collected per sample. Data was analyzed using Flowjo v10.10. A representative gating strategy can be found in Supporting Figure S1.

### mRNA production and purification

3xNLS-SpyCas9 mRNA used in this study was synthesized by in vitro transcription (IVT) using HiScribe T7 high-yield RNA synthesis kit (NEB, E2040S). CleanCap AG (3’OMe) and N1-Methyl-Pseudouridine-5’-Triphosphate were obtained from TriLink (N-7413-1 and N-1081-1). Following the manufacturer’s protocol, IVT reaction mixture was incubated at 37 °C for 2 hours, followed by DNase I treatment (NEB, M0303S) for 15 min to remove the DNA template. The resulting mRNA was purified using the Monarch Cleanup kit (NEB, T2050) and analyzed on a 1% glyoxal agarose gel (TriLink protocol). mRNA concentration was measured using Nanodrop. For *in vivo* studies, additional purification was performed by reverse-phase HPLC following the published protocol (33,36).

### LNPs formulation

LNPs were formulated with an amine-to-RNA-phosphate (N:P) ratio of 6:1. Lipid components were dissolved in 100% ethanol and combined at a total lipid concentration of 12.5 mM with the following molar ratios: 50 mol% 306-O12B, 10 mol% DOPC, 38.5 mol% Cholesterol, and 1.5 mol% DMG-PEG12k.

The RNA cargo was prepared at a 6:1 (w/w) ratio of SpyCas9 mRNA to sgRNA in a 10 mM citrate buffer prior to LNP assembly with a final RNA concentration of ∼0.35 mg/mL. LNPs were formed by microfluidic mixing of lipid and RNA solutions using the NanoAssemblr Ignite Nanoparticle Formulation System (Cytiva, 1001413) with a flow rate ratio of 3:1 (RNA:lipid) and a total flow rate of 12 mL/min.

Following formulation, LNPs were diluted 1:1 in PBS and subjected to buffer exchange in PBS overnight at 4°C with gentle stirring using 20K Slide-A-Lyzer Dialysis Cassettes (ThermoFisher, 66012). Dialyzed LNPs were filtered through 0.22 μm sterile syringe filters and concentrated using 100kDa MWCO Amicon Ultra centrifugal filters (Millipore, UFC8100). The resulting LNPs were stored at 4 °C prior to LNP characterization. Encapsulation efficiencies were determined using Quant-it Ribogreen Reagent and RNA Assay kit (ThermoFisher, R11490). Particle size and charge were measured by dynamic light scattering using the Malvern Zetasizer Lab system.

### Retro-orbital (RO) injection and *in vivo* studies

Animal experiments were performed in accordance with animal care ethnics approval and guidelines of the University of Massachusetts Medical School Institutional Animal Care and Use Committee (IACUC). Mice were 6-10 weeks old at the time of the experiments and weighed a day before injection day. For RO injections, mice were anesthetized in an approved induction chamber with 2-3% isoflurane in oxygen at 1.5 L/min. LNP formulations were administered via the retro-orbital sinus at doses of 0.2 and 0.8 mg/kg for *mPcsk9* and 0.3 and 1mg/kg for *mTtr*. Animals were monitored for signs of pain or distress and returned to clean cages once recovered. Two weeks post-injection, mice were euthanized, and livers were harvested for genomic DNA extraction.

### Genomic DNA extraction from cultured cells and mouse tissues

For cultured cells, genomic DNA (gDNA) was extracted 48-72 h after electroporation. Cell culture media were aspirated, and cells were lysed using QuickExtract DNA Extraction Solution (Lucigen), in accordance with the manufacturer’s protocol.

For mouse liver tissues, samples were flash-frozen and ground before processing. Tissue aliquots were transferred to microcentrifuge tubes, homogenized, and gDNA was extracted using DNeasy Blood and Tissue Kit (Qiagen, 69506) as recommended by the manufacturer.

### Indel analysis by targeted amplicon deep sequencing

A ∼200-250 bp genomic region of interest was PCR-amplified using NEBNext Ultra II Q5 Master Mix (NEB, M0544) and Q5 and Q7 adapter oligonucleotides for 30 cycles. Primers used for sequencing are provided in the Supporting Table 4 (Excel file). The amplified samples (2 μL) were used as a template for a second PCR (10 cycles) to introduce Q5 and Q7 index sequences. Barcoded PCR products were pooled and gel-extracted using the Zymo Gel DNA Recovery Kit and DNA Clean & Concentrator kits (Zymo Research 11-301, 11-303). To remove residual salts and gel contaminants, samples were further purified using SPRIselect bead-based reagent (Beckman Coulter, B23317) according to the manufacturer’s protocol.

The purified library was quantified using Qubit 1 x dsDNA HS Assay kits (ThermoFisher Q32851). Sequencing of pooled amplicons was performed on an Illumina NextSeq 2000 Sequencing system (P1 300-cycles kit), MiniSeq System (300-cycles kit, FC-420-1004), or MiSeq i100 Plus System (5M 300-cycles kit) following the manufacturer’s protocols.

The sequencing data were demultiplexed using bcl2fastq2 (Illumina) with barcode mismatches set to zero. FASTQ files were aligned, and editing efficiencies were quantified using CRISPResso2 (37) in batch mode with the following flags: -w 10, -q 30, -plot_window_size 20, and -min_frequency_alleles_around_cut_to_plot 0.1. Indel frequency = (inserted reads + deletion reads) / all aligned reads x 100%.

## Author Contributions

KAV, HZ, NA, GD, NB, AM, KY, JA, AK, SAW, EJS and JKW designed guide sequences. KAV, GD, JL, DE, JS, DC, AS and KY synthesized phosphoramidites and oligonucleotides. SAM and CL produced Cas9 protein and mRNA. KAV, HZ, NA, NG, SAM, ZC, PL, KP, MBH, CL, KL, JMRB, AVA, NB, and AM tested the properties of modified guide sequences. AK, SAW, NA, EJS and JKW conceived of the study and supervised the work. KAV, HZ, EJS and JKW wrote the paper. KY, SAW, KAV, HZ, EJS and JKW edited the paper.

## Data Availability

All data are presented in the main text and the supplementary information, with raw data available from the corresponding authors upon request.

## Supplementary Information

Online Supporting Information includes Figures S1-S5 and Supporting Tables 1-4 (Excel file).

## References

1. Khvorova, A. and Watts, J.K. (2017) The chemical evolution of oligonucleotide therapies of clinical utility. Nature Biotechnology, 35, 238–248.

2. Egli, M. and Manoharan, M. (2023) Chemistry, structure and function of approved oligonucleotide therapeutics. Nucleic Acids Research, 51, 2529–2573.

3. Tang, Q. and Khvorova, A. (2024) RNAi-based drug design: considerations and future directions. Nature Reviews Drug Discovery, 23, 341–364.

4. Rahdar, M., McMahon, M.A., Prakash, T.P., Swayze, E.E., Bennett, C.F. and Cleveland, D.W. (2015) Synthetic CRISPR RNA-Cas9-guided genome editing in human cells. Proc Natl Acad Sci U S A, 112, E7110–7117.

5. Yin, H., Song, C.Q., Suresh, S., Wu, Q., Walsh, S., Rhym, L.H., Mintzer, E., Bolukbasi, M.F., Zhu, L.J., Kauffman, K. et al. (2017) Structure-guided chemical modification of guide RNA enables potent non-viral in vivo genome editing. Nature Biotechnology, 35, 1179–1187.

6. O’Reilly, D., Kartje, Z.J., Ageely, E.A., Malek-Adamian, E., Habibian, M., Schofield, A., Barkau, C.L., Rohilla, K.J., DeRossett, L.B., Weigle, A.T. et al. (2018) Extensive CRISPR RNA modification reveals chemical compatibility and structure-activity relationships for Cas9 biochemical activity. Nucleic Acids Research, 47, 546–558.

7. Mir, A., Alterman, J.F., Hassler, M.R., Debacker, A.J., Hudgens, E., Echeverria, D., Brodsky, M.H., Khvorova, A., Watts, J.K. and Sontheimer, E.J. (2018) Heavily and fully modified RNAs guide efficient SpyCas9-mediated genome editing. Nature Communications, 9, 2641.

8. Finn, J.D., Smith, A.R., Patel, M.C., Shaw, L., Youniss, M.R., van Heteren, J., Dirstine, T., Ciullo, C., Lescarbeau, R., Seitzer, J., et al. (2018) A Single Administration of CRISPR/Cas9 Lipid Nanoparticles Achieves Robust and Persistent Genome Editing. Cell Reports, 22, 2227–2235.

9. Wu, J. and Yin, H. (2019) Engineering guide RNA to reduce the off-target effects of CRISPR. Journal of Genetic and Genomics, 46, 523–529.

10. Hassler, M.R., Turanov, A.A., Alterman, J.F., Haraszti, R.A., Coles, A.H., Osborn, M.F., Echeverria, D., Nikan, M., Salomon, W.E., Roux, L. et al. (2018) Comparison of partially and fully chemically-modified siRNA in conjugate-mediated delivery in vivo. Nucleic Acids Research, 46, 2185–2196.

11. Kliuchnikov, E., Maksudov, F., Zuber, J., Hyde, S., Castoreno, A., Waldron, S., Schlegel, M.K., Marx, K.A., Maier, M.A. and Barsegov, V. (2025) Improving the potency prediction for chemically modified siRNAs through insights from molecular modeling of individual sequence positions. Molecular Therapy Nucleic Acids, 36.

12. Zhang, H., Kelly, K., Lee, J., Echeverria, D., Cooper, D., Panwala, R., Amrani, N., Chen, Z., Gaston, N., Wagh, A. et al. (2023) Self-delivering, chemically modified CRISPR RNAs for AAV co-delivery and genome editing in vivo. Nucleic Acids Research, 52, 977–997.

13. Hendel, A., Bak, R.O., Clark, J.T., Kennedy, A.B., Ryan, D.E., Roy, S., Steinfeld, I., Lunstad, B.D., Kaiser, R.J., Wilkens, A.B. et al. (2015) Chemically modified guide RNAs enhance CRISPR-Cas genome editing in human primary cells. Nature Biotechnology, 33, 985–989.

14. Ryan, D.E., Taussig, D., Steinfeld, I., Phadnis, S.M., Lunstad, B.D., Singh, M., Vuong, X., Okochi, K.D., McCaffrey, R., Olesiak, M. et al. (2017) Improving CRISPR–Cas specificity with chemical modifications in single-guide RNAs. Nucleic Acids Research, 46, 792–803.

15. Cromwell, C.R., Sung, K., Park, J., Krysler, A.R., Jovel, J., Kim, S.K. and Hubbard, B.P. (2018) Incorporation of bridged nucleic acids into CRISPR RNAs improves Cas9 endonuclease specificity. Nature Communications, 9, 1448.

16. Nishimasu, H., Ran, F.A., Hsu, P.D., Konermann, S., Shehata, S.I., Dohmae, N., Ishitani, R., Zhang, F. and Nureki, O. (2014) Crystal structure of Cas9 in complex with guide RNA and target DNA. Cell, 156, 935–949.

17. Jinek, M., Jiang, F., Taylor, D.W., Sternberg, S.H., Kaya, E., Ma, E., Anders, C., Hauer, M., Zhou, K., Lin, S. et al. (2014) Structures of Cas9 endonucleases reveal RNA-mediated conformational activation. Science, 343, 1247997.

18. Palermo, G. (2019) Structure and Dynamics of the CRISPR–Cas9 Catalytic Complex. Journal of Chemical Information and Modeling, 59, 2394–2406.

19. Saha, K., Sontheimer, E.J., Brooks, P.J., Dwinell, M.R., Gersbach, C.A., Liu, D.R., Murray, S.A., Tsai, S.Q., Wilson, R.C., Anderson, D.G. et al. (2021) The NIH Somatic Cell Genome Editing program. Nature, 592, 195–204.

20. Hougland, J.L. and Piccirilli, J.A. (2009), Methods in Enzymology. Academic Press, Vol. 468, pp. 107–125.

21. Pagratis, N.C., Bell, C., Chang, Y.-F., Jennings, S., Fitzwater, T., Jellinek, D. and Dang, C. (1997) Potent 2′-amino-, and 2′-fluoro-2′-deoxyribonucleotide RNA inhibitors of keratinocyte growth factor. Nature Biotechnology, 15, 68–73.

22. Hoshika, S., Minakawa, N. and Matsuda, A. (2004) Synthesis and physical and physiological properties of 4’-thioRNA: application to post-modification of RNA aptamer toward NF-kappaB. Nucleic Acids Research, 32, 3815–3825.

23. Haeberli, P., Berger, I., Pallan, P.S. and Egli, M. (2005) Syntheses of 4’-thioribonucleosides and thermodynamic stability and crystal structure of RNA oligomers with incorporated 4’-thiocytosine. Nucleic Acids Research, 33, 3965–3975.

24. Yamada, K., Hariharan, V.N., Caiazzi, J., Miller, R., Ferguson, C.M., Sapp, E., Fakih, H.H., Tang, Q., Yamada, N., Furgal, R.C. et al. (2025) Enhancing siRNA efficacy in vivo with extended nucleic acid backbones. Nature Biotechnology, 43, 904–913.

25. Pater, A.A., Barber, H.M., Sudhakar, S., Chilamkurthy, R., Jana, S.K., Parasrampuria, M.A., Bosmeny, M.S., Graczyk-Marrs, J.A., Eddington, S.B., Blazier, C.A. et al. (2026) Chemical control of 2’-hydroxyl-dependent Cas9 target engagement enables CRISPR RNA ribose replacement. bioRxiv, 2026.2001.2026.701763.

26. Certo, M.T., Ryu, B.Y., Annis, J.E., Garibov, M., Jarjour, J., Rawlings, D.J. and Scharenberg, A.M. (2011) Tracking genome engineering outcome at individual DNA breakpoints. Nature Methods, 8, 671–676.

27. Iyer, S., Mir, A., Vega-Badillo, J., Roscoe, B.P., Ibraheim, R., Zhu, L.J., Lee, J., Liu, P., Luk, K., Mintzer, E. et al. (2022) Efficient Homology-Directed Repair with Circular Single-Stranded DNA Donors. CRISPR J, 5, 685–701.

28. Chen, Z., Kwan, S.Y., Mir, A., Hazeltine, M., Shin, M., Liang, S.Q., Chan, I.L., Kelly, K., Ghanta, K.S., Gaston, N., et al. (2023) A Fluorescent Reporter Mouse for In Vivo Assessment of Genome Editing with Diverse Cas Nucleases and Prime Editors. CRISPR J, 6, 570–582.

29. Hyjek-Składanowska, M., Vickers, T.A., Napiórkowska, A., Anderson, B.A., Tanowitz, M., Crooke, S.T., Liang, X.-h., Seth, P.P. and Nowotny, M. (2020) Origins of the Increased Affinity of Phosphorothioate-Modified Therapeutic Nucleic Acids for Proteins. Journal of the American Chemical Society, 142, 7456–7468.

30. Frey, P.A. and Sammons, R.D. (1985) Bond Order and Charge Localization in Nucleoside Phosphorothioates. Science, 228, 541–545.

31. Crooke, S.T., Vickers, T.A. and Liang, X.H. (2020) Phosphorothioate modified oligonucleotide-protein interactions. Nucleic Acids Research, 48, 5235–5253.

32. Chen, S., Heendeniya, S.N., Le, B.T., Rahimizadeh, K., Rabiee, N., Zahra, Q.U.A. and Veedu, R.N. (2024) Splice-Modulating Antisense Oligonucleotides as Therapeutics for Inherited Metabolic Diseases. BioDrugs, 38, 177–203.

33. Chen, Z., Kelly, K., Cheng, H., Dong, X., Hedger, A.K., Li, L., Sontheimer, E.J. and Watts, J.K. (2023) In Vivo Prime Editing by Lipid Nanoparticle Co-Delivery of Chemically Modified pegRNA and Prime Editor mRNA. GEN Biotechnology, 2, 490–502.

34. Birkenshaw, A., Thomson, T., Truong, M.P., Komaki, Y., Ramsden, N., Timpano, A., Huang, C., Blakney, A.K., Kurek, D.Z., Kulkarni, J. and Ross, C.J.D. (2026) sgRNA amount is a limiting factor in adenine base editing using RNA LNPs. Molecular Therapy Nucleic Acids, 37.

35. Wu, Y., Zeng, J., Roscoe, B.P., Liu, P., Yao, Q., Lazzarotto, C.R., Clement, K., Cole, M.A., Luk, K., Baricordi, C. et al. (2019) Highly efficient therapeutic gene editing of human hematopoietic stem cells. Nat Med, 25, 776–783.

36. Weissman, D., Pardi, N., Muramatsu, H. and Karikó, K. (2013) HPLC purification of in vitro transcribed long RNA. Methods in Molecular Biology, 969, 43–54.

37. Clement, K., Rees, H., Canver, M.C., Gehrke, J.M., Farouni, R., Hsu, J.Y., Cole, M.A., Liu, D.R., Joung, J.K., Bauer, D.E. and Pinello, L. (2019) CRISPResso2 provides accurate and rapid genome editing sequence analysis. Nature Biotechnology, 37, 224–226.

