## Supplementary Material for "Fully Modified SpyCas9 Guide RNAs Enable Robust Genome Editing In Cells and In Vivo"

**Contents:**

Supporting Figure S1. Flow gating strategy 2

Supporting Figure S2. Editing of TLR-MCV reporter by conjugated guides 3

Supporting Figure S3. Structural context of key tracrRNA positions within Cas9 4

Supporting Figure S4. Editing of mPcsk9 by conjugated guides 5

Supporting Figure S5. Gel analysis of sgRNA purity using Fragment Analyzer 6

Supporting Tables 1-4 containing all sequences and chemical modifications are separately attached (Excel file).

**Supporting Figure S1**

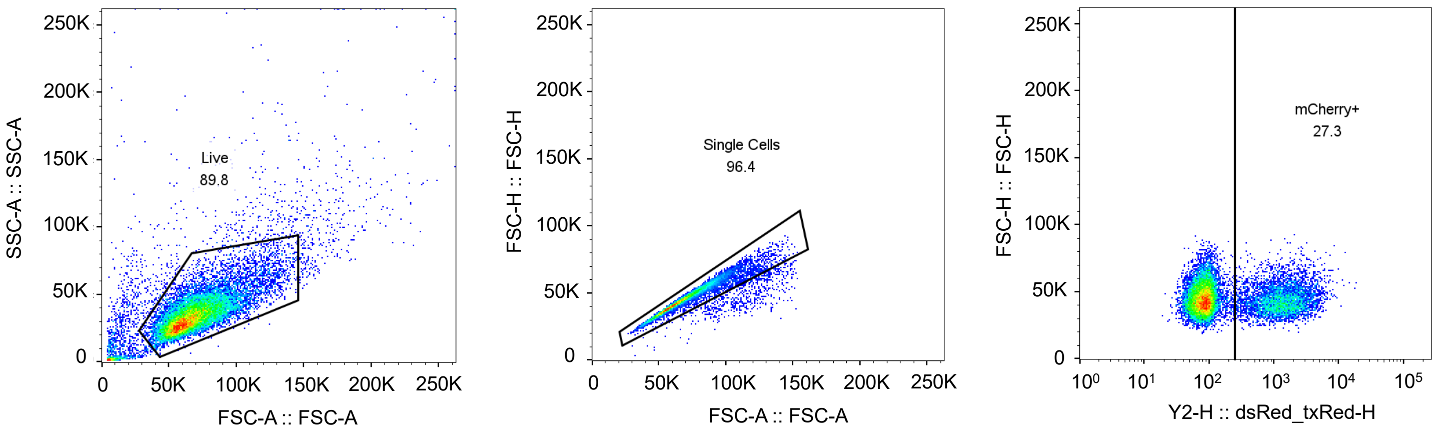


**Supporting Figure S1**. Representative flow cytometry gating strategy for HEK293T-TLR-MCV1 cell line.

**Supporting Figure S2**

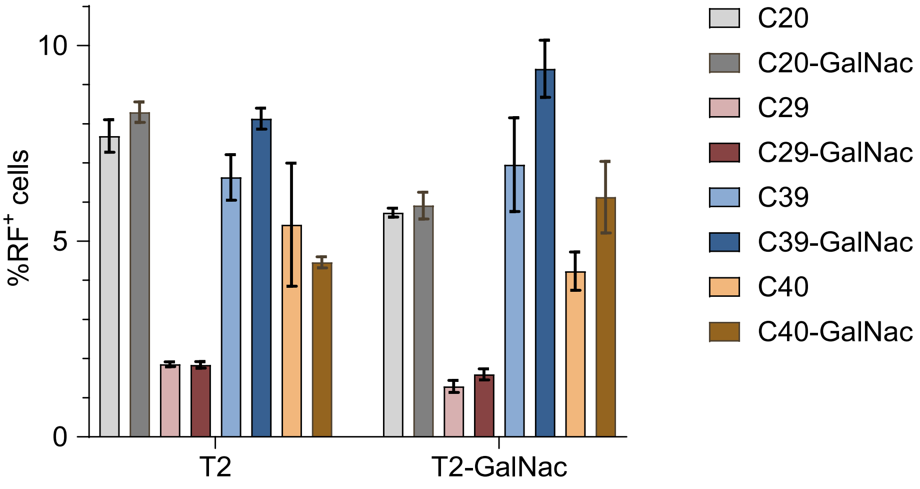


**Supporting Figure S2**. Editing efficiency in HEK293T-TLR cells following electroporation of 2 pmol RNPs targeting MCV1a, assembled using GalNac-conjugated or non-conjugated crRNAs and tracrRNAs. Editing efficiency was assessed by quantifying the percentage of red fluorescent-positive ($RF^+$) cells via flow cytometry. Data represent mean ± SD from two biological replicates.

**Supporting Figure S3**

**(A)**

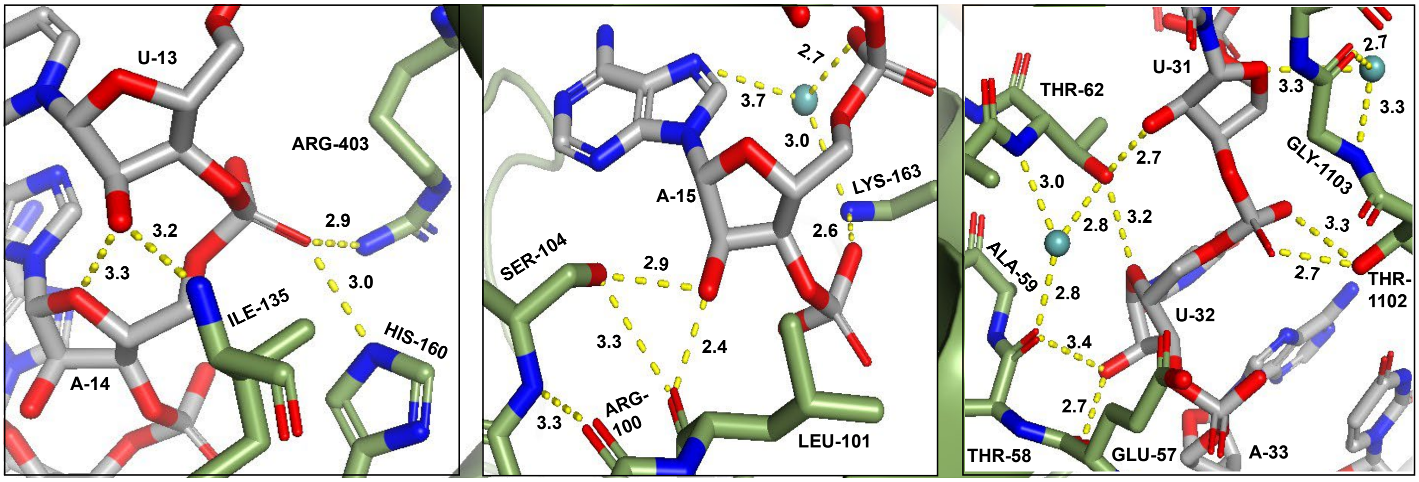


**(B)**

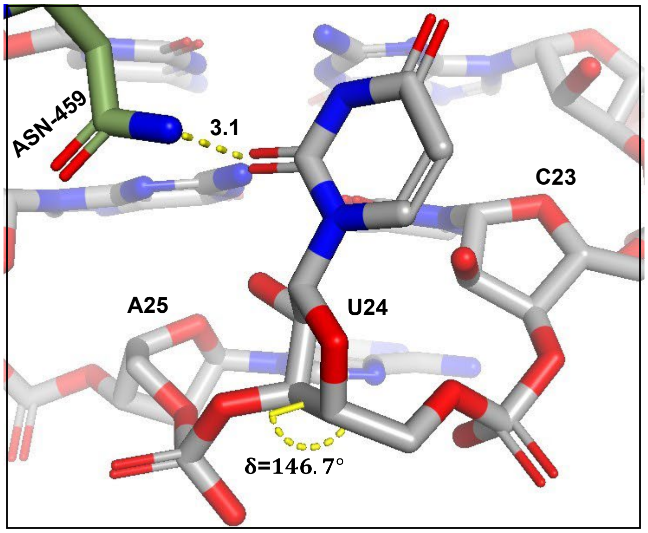


**Supporting Figure S3**. (**A**) Structure context of tracrRNA positions U13, A15, and U31-32 (corresponding to U45, U47, and U63-64 in the sgRNA, PDB ID: 4OO8) in the Cas9-sgRNA-DNA complex (15). (**B**) Structure context of tracrRNA positions C23 and U24 (corresponding to C55, U56 in 4OO8) in the Cas9-sgRNA-DNA complex (15). The sgRNA is shown in gray and SpyCas9 in green. $\boldsymbol{\delta}$ torsion angle (C5'-C4'-C3'-O3') at tracrRNA U24 is 146.7°, consistent with a C2'-endo sugar pucker.

**Supporting Figure S4**

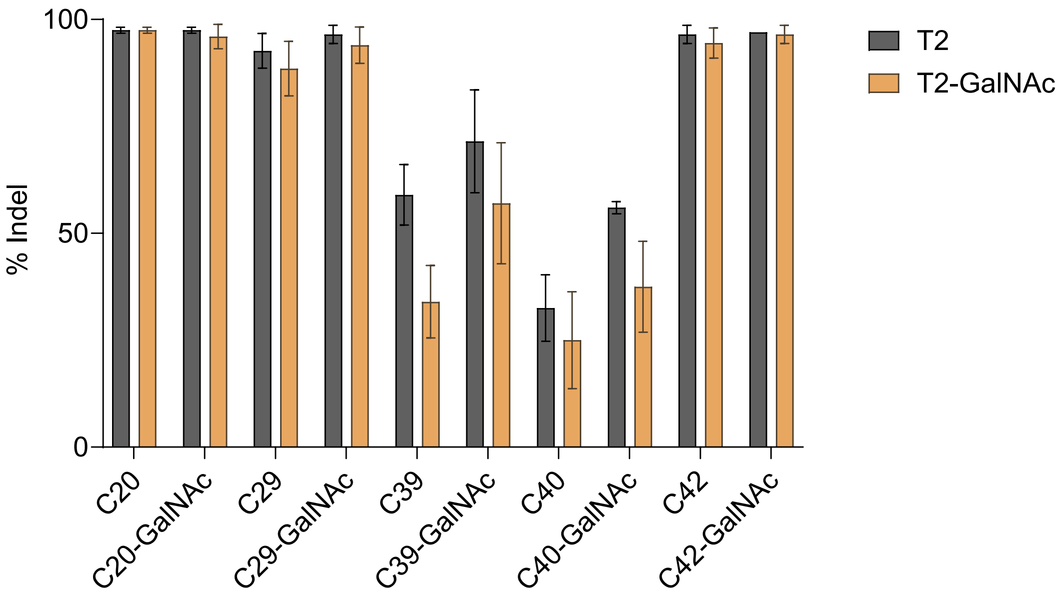


**Supporting Figure S4.** 3'-terminal conjugates are tolerated by heavily and fully modified crRNAs and tracrRNAs. Editing efficiency in Hepa1-6 cells following electroporation with 5 pmol RNPs targeting Pcsk9, assembled using **C20**, **C29**, **C39**, **C40**, and **C42** crRNAs and **T2** tracrRNA with or without GalNAc conjugated. Indel percentages were quantified by amplicon sequencing. Data represent mean ± SD from two biological replicates.

**Supporting Figure S5**

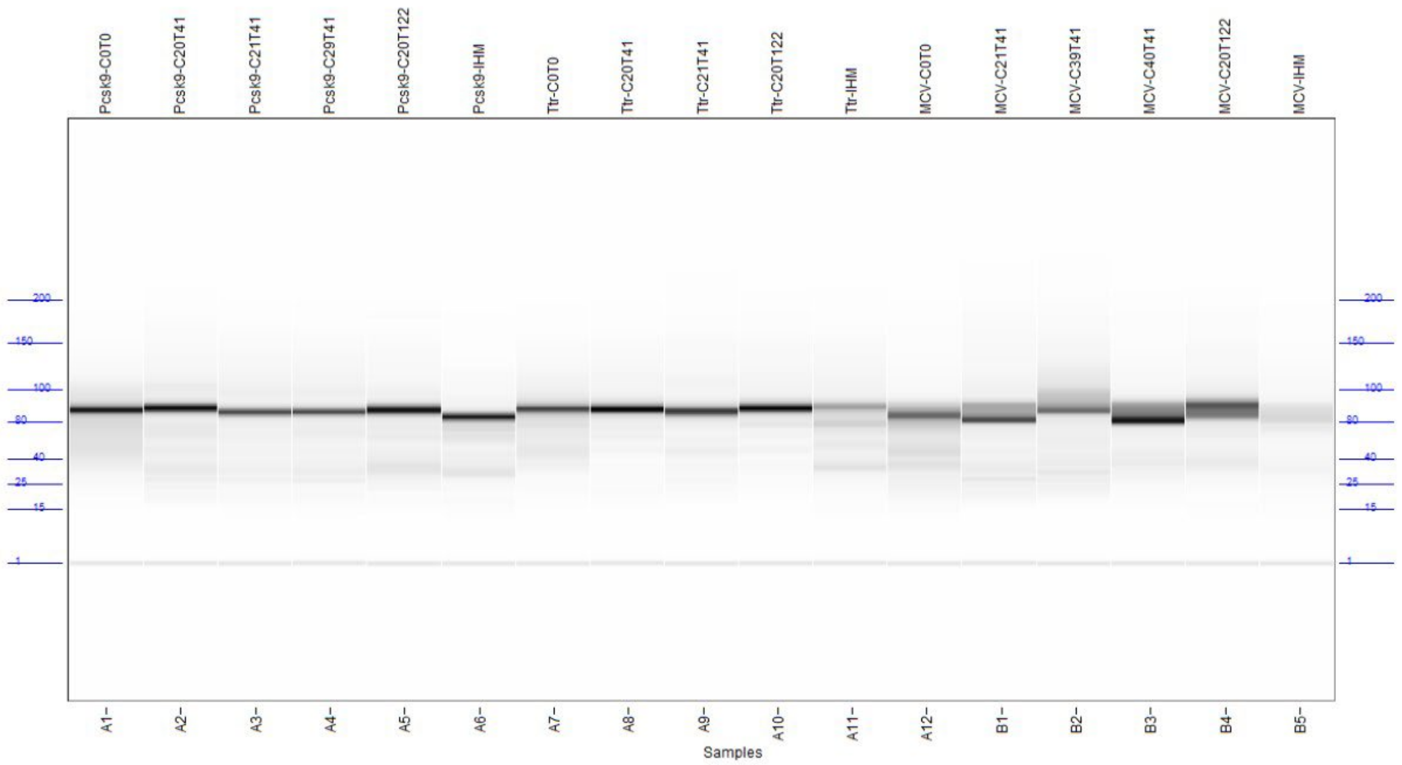


**Supporting Figure S5.** Gel image of sgRNAs used in this study generated using the Fragment Analyzer small RNA assay.
